# Tree-Aggregated Predictive Modeling of Microbiome Data

**DOI:** 10.1101/2020.09.01.277632

**Authors:** Jacob Bien, Xiaohan Yan, Léo Simpson, Christian L. Müller

## Abstract

Modern high-throughput sequencing technologies provide low-cost microbiome survey data across all habitats of life at unprecedented scale. At the most granular level, the primary data consist of sparse counts of amplicon sequence variants or operational taxonomic units that are associated with taxonomic and phylogenetic group information. In this contribution, we leverage the hierarchical structure of amplicon data and propose a data-driven and scalable tree-guided aggregation framework to associate microbial subcompositions with response variables of interest. The excess number of zero or low count measurements at the read level forces traditional microbiome data analysis workflows to remove rare sequencing variants or group them by a fixed taxonomic rank, such as genus or phylum, or by phylogenetic similarity. By contrast, our framework, which we call `trac` (`tr`ee-`a`ggregation of `c`ompositional data), learns data-adaptive taxon aggregation levels for predictive modeling, greatly reducing the need for user-defined aggregation in preprocessing while simultaneously integrating seamlessly into the compositional data analysis framework. We illustrate the versatility of our framework in the context of large-scale regression problems in human gut, soil, and marine microbial ecosystems. We posit that the inferred aggregation levels provide highly interpretable taxon groupings that can help microbiome researchers gain insights into the structure and functioning of the underlying ecosystem of interest.

## Introduction

Microbial communities populate all major environments on earth and significantly contribute to the total planetary biomass. Current estimates suggest that a typical human-associated microbiome consists of ~ 10^13^ bacteria [1] and that marine bacteria and protists contribute to as much as 70% of the total marine biomass [2]. Recent advances in modern targeted amplicon and metagenomic sequencing technologies provide a cost effective means to get a glimpse into the complexity of natural microbial communities, ranging from marine and soil to host-associated ecosystems [3, 4, 5]. However, relating these large-scale observational microbial sequencing surveys to the structure and functioning of microbial ecosystems and the environments they inhabit has remained a formidable scientific challenge.

Microbiome amplicon surveys typically comprise sparse read counts of marker gene sequences, such as 16S rRNA, 18S rRNA, or internal transcribed spacer (ITS) regions. At the most granular level, the data are summarized in count or relative abundance tables of operational taxonomic units (OTUs) at a prescribed sequence similarity level or denoised amplicon sequence variants (ASVs) [6]. The special nature of the marker genes enables taxonomic classification [7, 8, 9, 10] and phylogenetic tree estimation [11], thus allowing a natural hierarchical grouping of taxa. This grouping information plays an essential role in standard microbiome analysis workflows. For example, a typical amplicon data preprocessing step uses the grouping information for count aggregation where OTU or ASV counts are pooled together at a higher taxonomic rank (e.g., the genus level) or according to phylogenetic similarity [12, 13, 14, 15, 16]. This approach reduces the dimensionality of the data set and avoids dealing with the excess number of zero or low count measurements at the OTU or ASV level. In addition, rare sequence variants with incomplete taxonomic annotation are often simply removed from the sample.

This common practice of aggregating to a fixed taxonomic or phylogenetic level and then removing rare variants comes with several statistical and epistemological drawbacks. A major limitation of the fixed-level approach to aggregation is that it forces a tradeoff between, on the one hand, using low-level taxa that are too rare to be informative (requiring throwing out many of them) and, on the other hand, aggregating to taxa that are at such a high level in the tree that one has lost much of the granularity in the original data. Aggregation to a fixed level attempts to impose an unrealistic “one-size-fits-all” mentality onto a complex, highly diverse system with dynamics that likely vary appreciably across the range of species represented. A fundamental premise of this work is that the decision of how to aggregate should not be made globally across an entire microbiome data set *a priori* but rather be integrated into the particular statistical analysis being performed. Many factors, both biological and technical, contribute to the question of how one should aggregate: biological factors include the characteristics of the ecosystem under study and the nature of the scientific question; technical aspects include the abundance of different taxa, the available quality of the sequencing data—including sequencing technology, sample sequencing depth, and sample size—all of which may affect the ability to distinguish nearby taxa.

Another important factor when considering the practice of aggregating counts is that standard amplicon counts only carry relative (or “compositional”) information about the microbial abundances and thus require dedicated statistical treatment. When working with relative abundance data, the authors in [17, 18, 19] posit that counts should be combined with geometric averages rather than arithmetic averages. The common practice of performing arithmetic aggregation of read counts to some fixed level before switching over to the geometric-average-based compositional data analysis workflow is unsatisfactory since the “optimal” level for fixed aggregation is likely data-dependent, and the mixed use of different averaging operations complicates interpretation of the results.

To address these concerns, we propose a flexible, data-adaptive approach to tree-based aggregation that fully integrates aggregation into a statistical predictive model rather than relegating aggregation to preprocessing. Given a user-defined taxon *base level* (by default, the OTU/ASV level), our method `trac` (`tre`e-`a`ggregation of `c`ompositional data) learns dataset-specific taxon aggregation levels that are optimized for *predictive regression* modeling, thus making user-defined aggregation obsolete. Using OTU/ASVs as *base level*, Figure 1A illustrates the typical aggregation-to-genus level approach whereas Figure 1B shows the prediction-dependent `trac` approach. The `trac` method is designed to mesh seamlessly with the compositional data analysis framework by combining log-contrast regression [20] with tree-guided regularization, recently put forward in [21]. Thanks to the convexity of the underlying penalized estimation problem, `trac` can deliver interpretable aggregated solutions to large-scale microbiome regression problems in a fast and reproducible manner.

**Figure 1:**
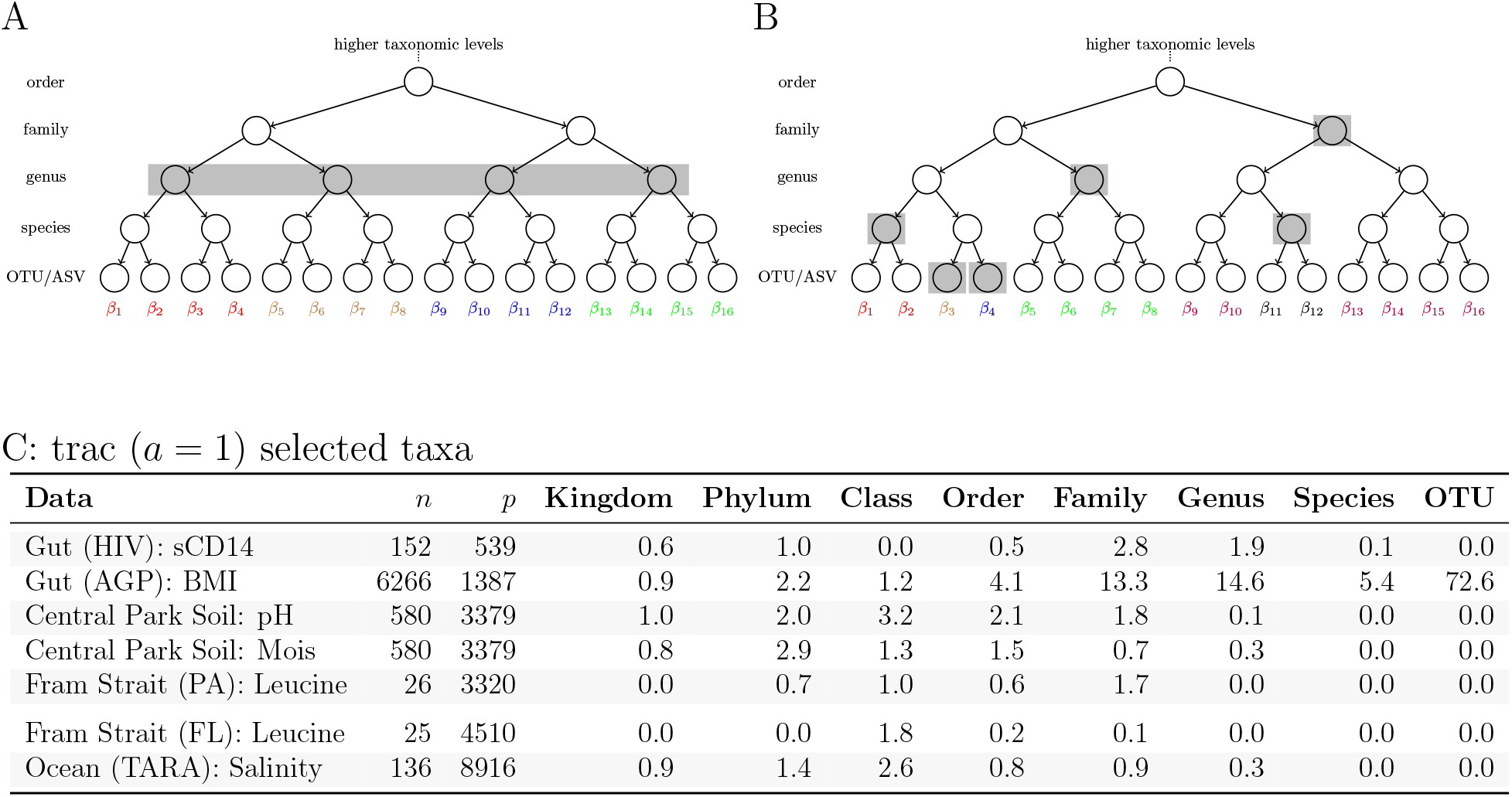
Illustration of fixed level and `trac`-based taxon aggregation. The trees represent the available taxonomic grouping of 16 *base level* taxa at the leaves (here OTU or ASV). A: Arithmetic aggregation of OTUs/ASVs to a fixed level (genus rank). All taxon base level counts are summed up to the respective parent genus. B: `trac`’s flexible tree-based aggregation in which the choice of what level to aggregate to can vary across the tree (e.g., two OTUs/ASVs, two species, one genus, and one family). The aggregation is based on the *geometric* mean of OTU/ASV counts and determined in a data-adaptive fashion with the goal of optimizing to the particular prediction task. C: Summary statistics of standard `trac`-inferred aggregation levels on all seven regression tasks. The Data column denotes the respective regression scenario (study name and outcome of interest), *n* the number of samples, and *p* the number of *base level* taxa (OTUs) in the data. The values in the taxonomic rank columns (Kingdom, Phylum, etc.) indicate the average number of taxa selected on that level by `trac` in the respective regression task. Averages are taken over ten random training/out-of-sample test data splits.

We demonstrate the versatility of our framework by analyzing seven representative regression problems on five datasets covering human gut, soil, and marine microbial ecosystems. Figure 1C summarizes the seven scenarios in terms of size of the microbial datasets and the average number of taxonomic aggregation levels selected by `trac`-inferred in the respective regression tasks. For instance, for the prediction of sCD14 concentrations (an immune marker in HIV patients) from gut microbiome data, `trac` selects, on average (over ten random training/test experiments), more taxa at the family level than any other taxonomic level, while it selects no taxa at the class or OTU level. By contrast, for the prediction of pH in the Central Park Soil data, class level taxa are selected more on average than any other level. This highlights the considerable departure from a typical fixed-level aggregation when prediction is the goal. Furthermore, the variability across the seven scenarios suggests that different amounts of aggregation may be warranted in different data sets.

Our `trac` framework complements other statistical approaches that make use of the available taxonomic or phylogenetic structure in microbial data analysis. For example, [22] uses phylogenetic information in the popular `unifrac` metric to measure distances between microbial compositions. The authors in [23, 24, 25, 26] combine tree information with the idea of “balances” from compositional data analysis [18] to perform phylogenetically-guided factorization of microbiome data. Others have included the tree structure in linear mixed models [27, 28], use phylogenetic-tree-based regression for detecting evolutionary shifts in trait evolution [29], and integrate tree-information into regression models for microbiome data [30, 31].

Along with our novel statistical formulation, we offer an easy-to-use and highly scalable software framework for simultaneous taxon aggregation and regression, available in the R package `trac` at https://github.com/jacobbien/trac. The R package `trac` also includes a fast solver for standard sparse log-contrast regression [15] to facilitate comparative analyses and a comprehensive documentation and workflow vignette. All data and scripts to fully reproduce the results in this manuscript are available on Zenodo at https://doi.org/10.5281/zenodo.4734527.

We next introduce `trac`’s mathematical formulation and discuss the key statistical and computational components of the framework. We also give an overview of the microbial data set collection and the comparative benchmark scenarios. To give a succinct summary of the key aspects of `trac` modeling on microbiome data, we will present and discuss three of the seven regression scenarios in detail. The other scenarios are available in the Supplementary Material. We conclude the study by highlighting key observations and provide recommendations and viable extensions of the `trac` framework.

## Materials and methods

### Modeling strategy

Let *y* ∈ ℝ^*n*^ be *n* observations of a variable we wish to predict and let 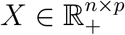 be a matrix with *X_ij_* giving the number of (amplicon) reads assigned to taxon *j* in sample *i*. The total number of reads ∑_*j*_ *X_ij_* in sample *i* is a reflection of the sequencing process and therefore should not be interpreted as providing meaningful information about the biological sample itself. This observation has motivated the adoption of compositional data methods, which ensure that analyses depend only on *relative* abundances. Following the foundational work in [20], one appropriate model for regression with relative abundance data is the log-contrast model where the outcome of interest is modeled as linear combinations of log-ratios (i.e., log-contrasts) of relative abundance features. For high-dimensional microbiome data, the authors in [15] propose solving an ℓ_1_-penalized regression estimator that includes a zero-sum constraint on the coefficients, the so-called sparse log-contrast model. Writing log(*X*) for the matrix with *ij*th entry log(*X_ij_*), their estimator is of the form

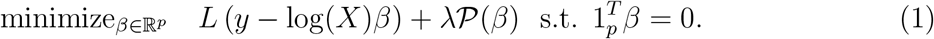

Here, *L*(*r*) = (2*n*)^−1^ ||*r*||^2^ is the squared error loss and 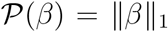 is the ℓ_1_ penalty [32]. The zero-sum constraint ensures that this model is equivalent to a log-contrast model [33] and invariant to sample-specific scaling. To understand the intuition behind the sparse log-contrast model, imagine that *β_j_* and *β_k_* are the only two nonzero coefficients. In such a case, the zero-sum constraint implies that predictions will be based on only the log-ratio of these two taxa. This can be seen by noting that *β_j_* = –*β_k_*, and so our model’s prediction for observation *i* would be given by the following:

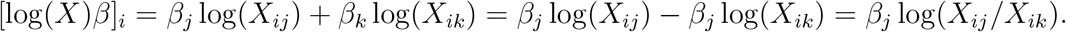

Thus, using a log has the effect of turning differences into ratios. In addition, the zero-sum constraint provides invariance to sample-specific scaling: Replacing *X* by *DX*, where *D* is an arbitrary diagonal matrix, leaves Eq. (1) unchanged:

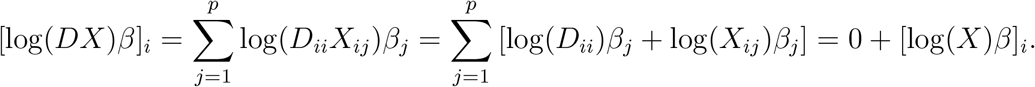

The choice of the ℓ_1_ penalty was motivated in [15] by the high dimensionality of microbiome data and the desire for parsimonious predictive models. However, such a penalty is not well-suited to situations in which large numbers of features are highly rare [21], a well-known feature of amplicon data. A common remedy, also adopted in [15], is to aggregate taxa at the base level, e.g., OTUs or ASVs, to the genus level and then to screen out all but the most abundant genera. Figure 1A depicts this standard practice: taxonomic (or phylogenetic) information in the form of a tree 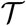 is used to aggregate data, usually in an arithmetic manner (i.e. by summing), to a *fixed level* of the tree.

Our goal is to make aggregation more flexible (as illustrated in Figure 1B), to allow the prediction task to inform the decision of how to aggregate, and to do so in a manner that is consistent with the log-contrast framework introduced above. A key insight is that aggregating features can be equivalently expressed as setting elements of *β* equal to each other. For example, suppose we partition the *p* base level taxa into *K* groups *G*_1_,…, *G_K_* and demand that *β* be constant within each group. Doing so yields K aggregated features. If all of the *β_j_* in group *G_k_* are equal to some common value, then

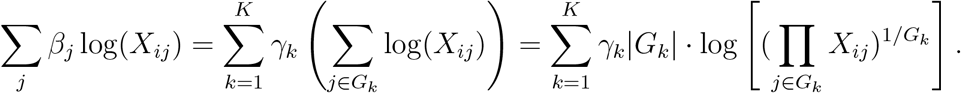

Thus, we are left with a linear model with *K* aggregated features, each being proportional to the log of the geometric mean of the base level taxa counts.

Associating the elements of *β* with the leaves of 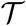, the above insight tells us that if our estimate of *β* is constant within subtrees of 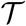, then that corresponds to a regression model with tree-aggregated features. In particular, each subtree with constant *β*-values will correspond to a feature, which is the log of the geometric mean of the counts within that subtree. The `trac` estimator uses a convex, tree-based penalty 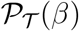 for the penalty in Eq. (1) that is specially designed to promote *β* to have this structure that is based on subtrees of 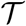. The mathematical form of 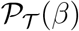 is given in Supplementary Material B. There, we show that the `trac` estimator reduces to solving the optimization problem:

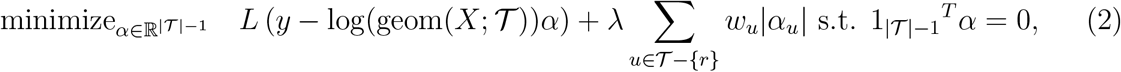

where 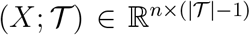 is a matrix where each column corresponds to a non-root node of 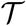 and consists of the geometric mean of all base level taxa counts within the subtree rooted at *u*. Comparing this form of the `trac` optimization problem to Eq. (1) reveals an alternate perspective: `trac` can be interpreted as being like a sparse log-contrast model but instead of the features corresponding to base level taxa, they correspond to the geometric means of non-root taxa in 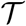 (i.e., *X* is replaced by 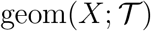). This also facilitates model interpretability since we can directly combine positive and negative predictors into pairs of log-ratio predictors. For example, if taxa *α_u_* > 0 and *α_v_* < 0 are the only nonzero coefficients, then our predictions would be based on

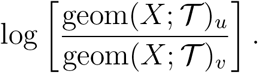

The particular choice of penalty is a weighted ℓ_1_-norm. While the `trac` package allows the user to specify general choices of weights *w_u_* > 0, a convenient and interpretable strategy is to set weights to be an inverse power of the number of leaves in the subtree rooted at *u, w_u_* = |*L_u_*|^−*a*^. The scalar parameter *a* ∈ ℝ controls the overall aggregation strength, with *a* =1 being the default setting in `trac`. If the user decreases a, `trac` favors aggregations at a lower level of the tree. For *a* sufficiently negative, `trac` admits solutions equivalent to a sparse log-contrast model without aggregation since only leaves (with |*L_u_*| = 1) will remain unaffected by the weight scaling. The regularization parameter λ, on the other hand, is a positive number determining the overall tradeoff between prediction error on the training data and how much aggregation should occur. By varying λ, we can trace out an entire solution path 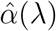, from highly sparse solutions (large λ) to more dense solutions involving many taxa (small λ). This “aggregation path” can itself be a useful exploratory tool in that it provides an ordering of the taxa as they enter the model.

### Computation, model selection, and prediction

Using `trac` in practice requires the efficient and accurate numerical solution of the convex optimization problem, specified in Eq. (2), across the full aggregation path. We experimented with several numerical schemes and found the path algorithm of [34] particularly well-suited for this task. The `trac` R package internally uses the path algorithm implementation from the `c-lasso` Python package [35], efficiently solving even high-dimensional `trac` problems. The `trac` package also provides a fast implementation of sparse log-contrast regression [15] for model comparison. The R package `reticulate` [36] is instrumental in connecting `trac` with the underlying Python library. The R packages `phyloseq` [37], `ggplot2` [38], `ape` [39], `igraph` [40], and `ggtree` [41] are used for operations on tree structures and visualization.

To find a suitable aggregation level along the solution path, we use cross validation (CV) with mean squared error to select the regularization parameter λ ∈ [λ_min_, λ_max_] for all the results presented in this paper. In particular, we perform 5-fold CV with the “one-standard-error rule” (1SE) [42], which identifies the largest λ whose CV error is within one standard error of the minimum CV error. This heuristic purposely favors models that involve fewer taxa and are therefore easier to interpret. (We also use the 1SE rule to select λ for the sparse log-contrast model.) The parameter a is a user-defined control parameter and not subject to a model selection criterion. Having solved the `trac` optimization problem and chosen a value of the tuning parameter 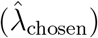, we can predict the response value at a new sample. Given a new vector of abundances 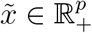, we predict the response to be

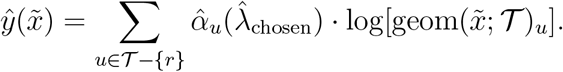

Due to `trac`’s sparsity penalty, in general only a small number of coefficients will be non-zero, and thus the predictions will depend on only a small number of taxas’ geometric means.

### Data collection

We assembled a collection of five publicly available and previously analyzed datasets, spanning human gut, soil, and marine ecosystems (see also Data column in Figure 1C). All datasets, except for Tara, consist of 16S rRNA amplicon data of Bacteria and Archaea in the form of OTU count tables, taxonomic classifications, and measured covariates, as provided in the original publications. For ease of interpretability, we leverage the taxonomic tree information rather than phylogeny in our aggregation framework. To investigate potential human host-microbiome interactions, we re-analyze two human gut datasets, one cohort of HIV patients (Gut [HIV]), available in [43], comprising *p* = 539 OTUs and *n* = 152 samples, and the other a subset of the American Gut Project data (Gut (AGP)) [5], provided in [44], comprising *p* = 1387 OTUs present in at least 10% of the *n* = 6266 samples. To study niche partitioning in terrestrial ecosystems, we use the Central Park soil dataset [45], as provided by [23], which consists of *p* = 3379 OTUs and *n* = 580 samples with a wide range of soil property measurements. For marine microbial ecosystems, we consider a sample collection from the Fram Strait in the North Atlantic [46], available at https://github.com/edfadeev/Bact-comm-PS85. The data set consists of *n* = 26 samples for *p* = 3320 OTUs in the particle-associated size class, and *n* = 25 samples for *p* = 4510 OTUs in the free-living size class. The second marine dataset is the Tara global surface ocean water sample collection [3], available at http://ocean-microbiome.embl.de/companion.html, which comprises metagenome-derived OTUs (mOTUs). In Tara, each of the *p* = 8916 mOTUs considered here is present in at least 10% of the *n* = 136 samples. All data and analysis scripts are available in fully reproducible R workflows at https://github.com/jacobbien/trac-reproducible. Since `trac` can operate on any taxon base level, we provide all data sets both in the form of the original (m)OTU base level as well as in arithmetically aggregated form on higher-order ranks, i.e., species, genus, family, order, class, and phylum. This facilitates straightforward method comparison across different base level aggregations.

### Method comparison and model quality assessment

To provide a comprehensive model performance evaluation and to highlight the flexibility of the `trac` modeling framework, we consider the following benchmark scenarios. Firstly, we consider three different regression models. We choose the sparse log-contrast regression model [15] as the standard baseline of performing regression on compositional data and can be considered as a limiting case of `trac`. In addition, we consider `trac` with two different aggregation parameters *a*. The setting *a* = 1 is referred to as standard `trac`. The setting *a* = 1/2 is referred to as *weighted* `trac` and tends to favor aggregations closer to the leaf level. Secondly, to assess the influence of arithmetic aggregation to a fixed level, e.g., the genus level, we compare the performance of all regression models for three different input base levels: OTU, genus, and family level.

To assess how well a log-contrast or `trac` model generalizes to “unseen” data, we randomly select 2*/*3 of the samples in each of the considered datasets for model training and selection. On the remaining 1/3 of the samples, we compute out-of-sample test mean squared error as well as the Pearson correlation between model predictions and actual measurements on the test set. While the out-of-sample test error serves as a key quantity to assess model generalizability, we also record overall model sparsity, measured in terms of number of aggregations (or taxa for sparse log-contrast models) in the trained model. Model sparsity serves as measure how “interpretable” a model is. Finally, we repeat all analysis on ten random training/test splits of the data to measure average test error and model sparsity. To ease interpretability, we analyze the trained models derived from split 1 in greater detail throughout the next section and detail the biological significance of the derived regression models.

## Results and Discussion

We next highlight key results of the `trac` framework for three of the seven regression scenarios described above on three different microbiome datasets. The first scenario considers the prediction of an immune marker (soluble sCD14) in HIV patients from microbiome data. In this scenario, we detail the behavior of a typical `trac` aggregation path and the model selection process. Furthermore, we compare the performance of `trac` models at different taxon base levels (OTU, genus, and family level) and aggregation weights (*a* = {1/2,1}) with standard sparse log-contrast models and analyze the resulting taxa aggregations. In the second scenario, we apply `trac` to predict pH concentrations in Central Park soil from microbial abundances and compare the resulting aggregations to known associations of pH and microbial taxa. The last scenario considers salinity prediction in the global ocean from Tara mOTU data. Further `trac` prediction scenarios are available in the Supplementary Material, including Body Mass Index (BMI) predictions on the American Gut Project Data, soil moisture prediction in Central Park soil, and primary productivity prediction from marine microbes in two different size fractions in the North Atlantic Fram Strait.

### Immune marker sCD14 prediction in HIV patients

Infection with HIV is often paired with additional acute or chronic inflammation events in the epithelial barrier, leading to disruption of intestinal function and the microbiome. The interplay between HIV infection and the gut microbiome has been posited to be a “two-way street” [47]: HIV-associated mucosal pathogenesis potentially leads to perturbation of the gut microbiome and, in turn, altered microbial compositions could result in ongoing disruption in intestinal homeostasis as well as secondary HIV-associated immune activation and inflammation.

Here, we investigate one aspect of this complex relationship by learning predictive models of immune markers from gut amplicon sequences. While [48] were among the first to provide evidence that gut microbial *diversity* is a predictor of HIV immune status (as measured by CD4+ cell counts), we consider soluble CD14 (sCD14) measurements in HIV patients as the variable to predict and learn an interpretable regression model from gut microbial amplicon data. sCD14 is a marker of microbial translocation and has been shown to be an independent predictor of mortality in HIV infection [49].

Following [43], we analyze a HIV cohort of *n* = 152 patients where sCD14 levels (in pg/ml units) and fecal 16S rRNA amplicon data were measured. Using as base level all available *p* = 539 bacterial and archaeal OTUs, we first illustrate the typical `trac` prediction and model selection outputs with default weight parameter *a* = 1 on the first (of overall ten) training/test splits in Figure 2. In Figure 2A, we visualize the solution of the *α* coefficients associated with each aggregation along the regularization path. The vertical lines indicate the aggregations that were selected via cross-validation (CV) with the Minimum Mean Squared Error (MSE, dotted line) and one-standard-error rule (1SE, dashed line) (see Figure 2B). On the test data, we highlight the relationship between test prediction performance of the `trac` models versus the number of inferred aggregations (Figure 2D). Models between five and 28 aggregations show excellent performance on the test set. `trac` with the 1SE rule identified a parsimonious model with aggregation to five main taxa (Figure 2E): the kingdom Bacteria, phylum Actinobacteria and the family Lachnospiraceae are negatively associated, and the family Ruminococcaceae and the genus Bacteroides are positively associated with sCD14 counts, thus resulting in a `trac` model with three log-contrasts.

**Figure 2:**
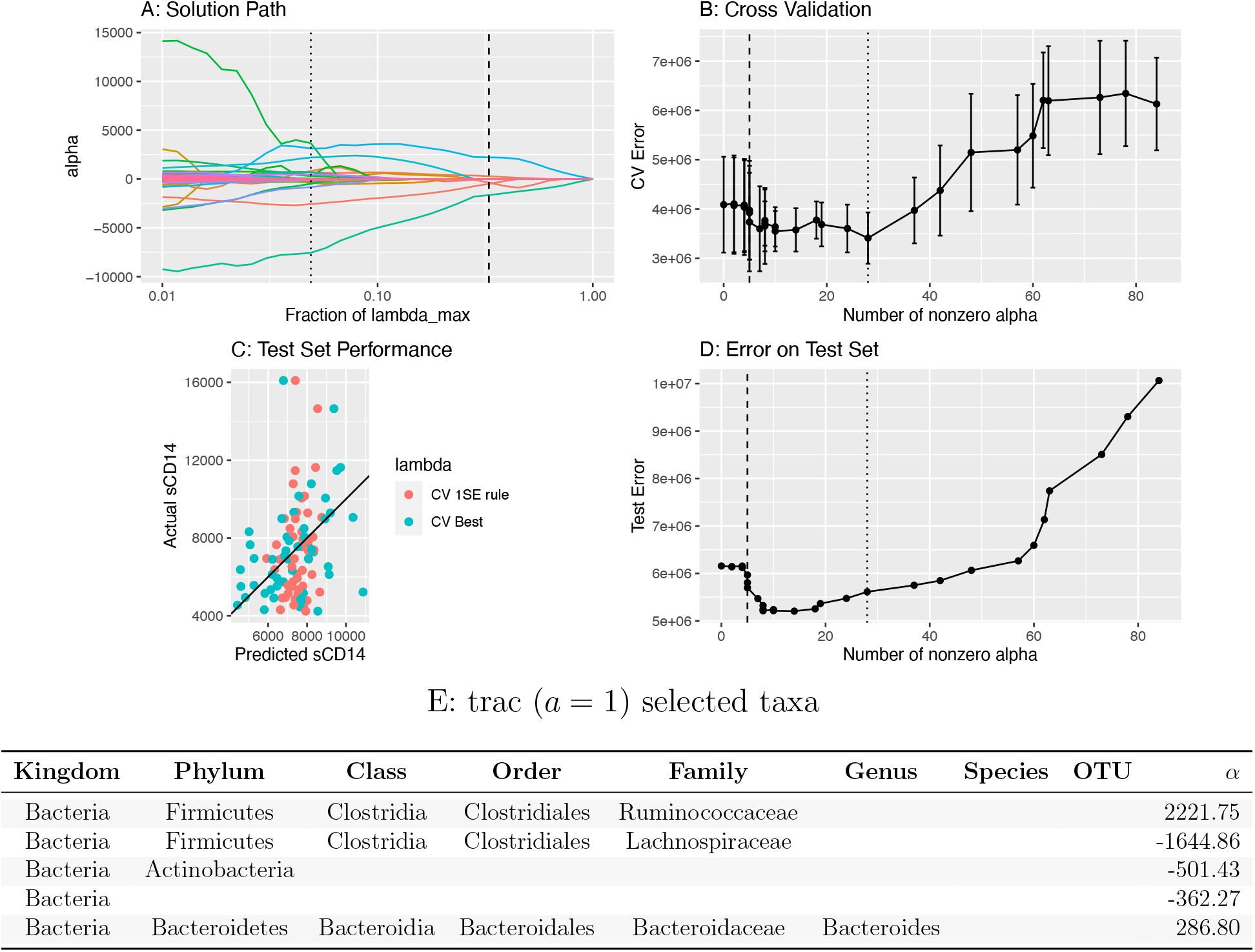
Overview of `trac` aggregation and model selection with standard weighting *a* =1 on the sCD14 data. A: Varying the `trac` regularization parameter λ produces a solution (aggregation) path. Each colored line corresponds to a distinct taxon, showing its *α* coefficient value as the tuning parameter λ increases. The larger λ is, the more coefficients are set to 0, leading to a more parsimonious model. The dotted and dashed vertical lines mark the λ-values selected by the CV best and 1SE rule, respectively. B: Illustration of the crossvalidation (CV) procedure. Mean (and standard error) CV error vs. λ path with selected λ values at best CV error (dotted vertical line) or with the 1SE rule (dashed vertical line) C: The actual vs. predicted values of sCD14 on the test set (1SE rule in red, CV best in blue). The Pearson correlation of `trac` predictions on the test set is 0.37 with the CV best solution and 0.23 with the CV 1SE rule, respectively. D: Error on the test set vs. number of selected aggregations. E: The `trac` model selected with the 1SE rule comprises five taxa across four levels, listed in the bottom table (see Figure 3A for tree visualization of the aggregations). The column labeled *α* gives the nonzero coefficient values, which are in the same units as the sCD14 response variable.

From a biological perspective, this `trac` analysis suggests a strong role of the Ruminococcaceae to Lachnospiraceae family ratio and, to a lesser extent, the Ruminococcaceae to Actinobacteria ratio in predicting mucosal disruption (as measured by sCD14). This follows from observing the large positive *α* coefficient associated with Ruminococcaceae and the large negative *α* coefficients associated with Lachnospiraceae and Actinobacteria (and recalling the interpretation of the `trac` output in terms of log-ratios). The protective or disruptive roles of Ruminococci or Lachnospiraceae in HIV patients is typically considered to be highly species-specific. Moreover, few consistent microbial patterns are known that generalize across studies [50]. For instance, [51] report high variability and diverging patterns of the differential abundances of individual OTUs belonging to the Ruminococcaceae and Lachnospiraceae family in HIV-negative and HIV-positive participants. Our model posits that, on the family level, consistent effects of these two families are detectable in amplicon data. This also suggests that, with the right aggregation level, a re-analysis of recent HIV-related microbiome data may, indeed, reveal reproducible patterns of different taxon groups in HIV infection.

To quantify the effect of taxon base level and aggregation weight scaling a, we re-analyze the data at OTU, genus, and family base level and compare `trac` models to sparse log-contrast models at the respective base level. The latter approach thus reflects the default mode of analysis, proposed in [15], where sparse log-contrast modeling on fixed genus aggregations was performed. Figure 3 visualizes the estimated `trac` aggregations (*a* = {1, 1/2}) and sparse taxa on the taxonomic tree of the sCD14 data.

**Figure 3:**
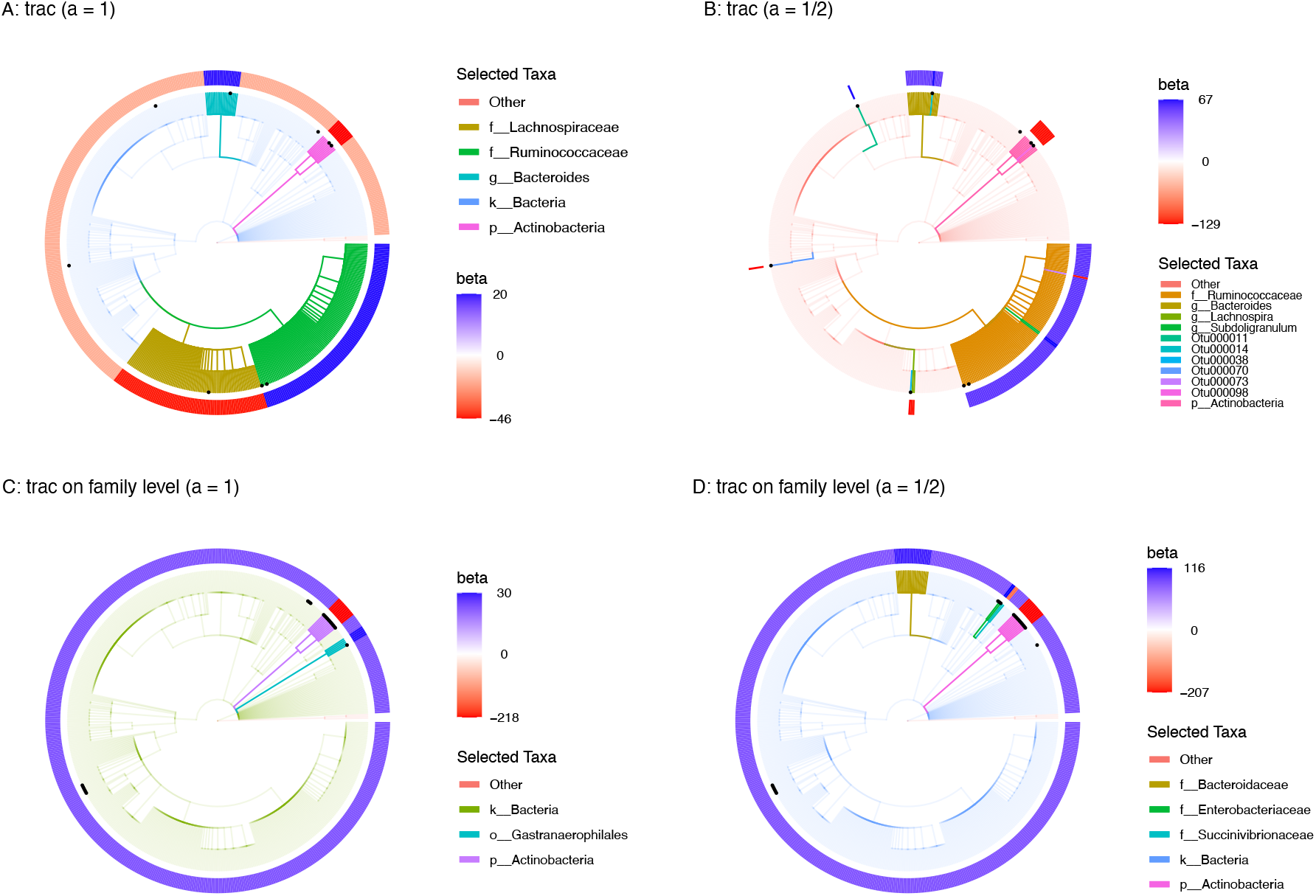
Taxonomic tree visualization of `trac` aggregations in four selected scenarios using sCD14 data (training/test split 1). Each tree represents the taxonomy of the *p* = 539 OTUs. Colored branches highlight the estimated `trac` taxon aggregations. The black dots mark the selected taxa of the respective sparse log-contrast model. The outer rim represents the value of *β* coefficients in the `trac` model from Eq. (1). A: Standard `trac` (*a* = 1) with OTUs as taxon base level selects five aggregations. B: Weighted `trac` (*a* = 1/2) with OTU base level selects eleven aggregations, including six on the OTU level. Four of these OTUs were also selected by the sparse log-contrast model which comprises nine OTUs in total (black dots) (see Suppl. Tables 6 and 7 for the selected coefficients). C: Standard `trac` (*a* = 1) with family base level selects three aggregations. D: Weighted `trac` (*a* = 1/2) with family as taxon base level selects five aggregations, including one family (Enterobacteriacaeae) shared with the sparse log-contrast model when also applied at the family base level (see Suppl. Tables 10 for the six selected families).

Figure 3A and B show the estimated models with OTUs as taxon base level, Figure 3C and D with family base level. Figure 3A highlights the previously discussed five aggregations from Figure 2E (Bacteria, Ruminococcaceae, Lachnospiraceae, Actinobacteria, and Bacteroides), found with standard `trac` (*a* = 1), by coloring the respective branches of the corresponding full taxonomic tree. We observe that the selected OTUs of the sparse log-contrast model (highlighted as black dots) cover each of the `trac` aggregations, including two OTUs in the phylum Actinobacteria, two OTUs in the family Ruminococcaceae, and one OTU in Lachnospiraceae family (see Suppl. Table 7 for the selected coefficients). Figure 3B highlights how weighted `trac` with *a* =1/2 results in predictive models that can represent a sort of compromise between both standard `trac` and sparse log-contrast components. For instance, weighted `trac` still comprises the Ruminococcaceae family, the Actinobacteria phylum, and the Bacteroides genus but also shares four OTUs with the sparse log-contrast model. This exemplifies the flexibility of the `trac` framework in fine-tuning predictive models to the “right” level of aggregation. We observe a similar but less pronounced effect of the weighting when using aggregated family counts as taxon base level (Figure 3C and D). The `trac` models comprise three and five aggregations, respectively, with the Actinobacteria phylum common to both. The sparse log-contrast model comprises six families, three of which are covered by the weighted `trac` model (two families in the Actinobacteria phylum and the Enterobacteriaceae family).

To compare the different statistical models in terms of interpretability and prediction quality, we report the sparsity level and the out-of-sample prediction errors, averaged over ten different training/test splits, in Table 1. We observe that for the sCD14 data set, standard `trac` with OTU base levels delivers the sparsest (on average, seven aggregations) and most predictive solution (average test error 6.3e+06), followed by standard `trac` on the family level (average test error 6.5e+06). The sparse log-contrast model with genus base level has considerably reduced prediction capability (average test error 7.1e+06). On this data set, weighted `trac` (*a* = 1/2) models show the expected intermediate properties between sparse log-contrast and standard `trac` solutions.

**Table 1:**
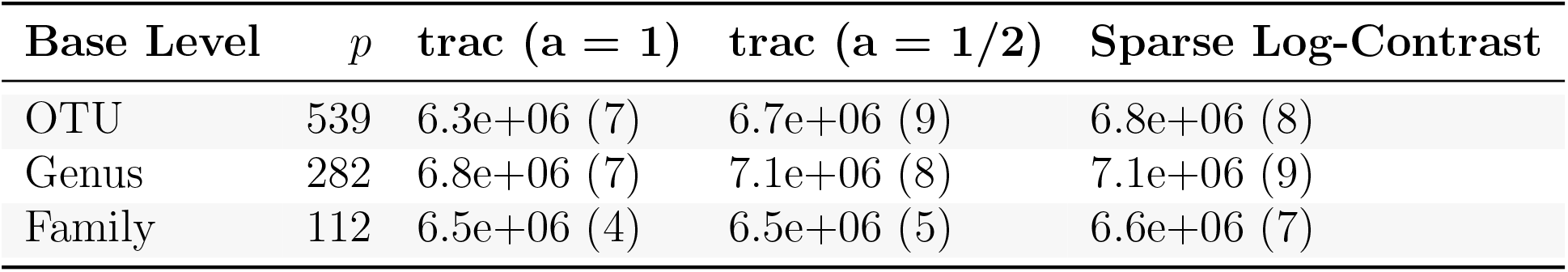
Average out-of-sample test errors (rounded average model sparsity in parenthesis) for `trac` (*a* = {1, 1/2}) and sparse log-contrast models, respectively. Each row considers a different base level (OTU, genus, and family). Each number is averaged over ten different training/test splits of the sCD14 data.

### Predicting Central Park soil pH concentration from microbiome data

We next perform `trac` prediction tasks on environmental rather than host-associated microbiome data. We first consider soil microbial compositions since they are known to vary considerably across spatial scales and are shaped by myriads of biotic and abiotic factors. Using univariate regression models, the authors in [52] found that soil habitat properties, in particular pH and soil moisture deficit (SMD), can predict overall microbial “phylotype” diversity. For instance, using *n* = 88 soil samples from North and South America, the authors in [53] showed that soil pH concentrations are strongly associated with amplicon sequence compositions, as measured by pairwise `unifrac` distances. Moreover, they found that soil pH correlated positively with the relative abundances of Actinobacteria and Bacteroidetes phyla, negatively with Acidobacteria, and not at all with Beta/Gammaproteobacteria ratios.

Here, we use `trac` on the Central Park soil data collection comprising *n* = 580 samples and *p* = 3379 bacterial and archaeal OTUs [45, 23] to provide a refined analysis of the relationship between soil microbiome and habitat properties. Rather than looking at the univariate correlative pattern between soil properties and phyla, we build multivariate models that take soil pH as the response variable of interest and optimize taxa aggregations using `trac` and sparse log-contrast models. The predictive analysis for soil moisture is relegated to the Supplementary Materials.

For pH prediction, standard `trac` gives an interpretable model with six aggregated taxonomic groups (see Figure 4A): the two phyla Bacteroidetes and Verrucomicrobia and the class Acidobacteria-6 were positively associated, whereas the order Acidobacteriales, the class Gammaproteobacteria, and the overall kingdom of Bacteria (compared to Archaea) were negatively associated with pH (see bottom table in Figure 4). We can thus associate a log-contrast model with three log-ratios of aggregated taxonomic groups with soil pH in Central Park. The overall Pearson correlation between the `trac` predictive model and the training data was 0.68. On the test data, the model still maintained a high correlation of 0.65. With the standard caveat that regression coefficients do not have the same interpretation (or even necessarily have the same sign) as their univariate counterparts, our model also supports a positive relationship between the Bacteroidetes phylum and pH and gives refined insights into the role of the Acidobacteria phylum. The model posits that the class Acidobacteria-6 is positively related and the order Acidobacteriales (in the Acidobacteriia class) is negatively related with pH. The authors in [23] observed similar groupings in their phylofactorization of the Central Park soil data. There, the classes Acidobacteria-6 and Acidobacteriia belonged to different “binned phylogenetic units” whose relative abundances increased and decreased along the pH gradient, respectively. Finally, the phylum Verrucomicrobia and the class Gammaproteobacteria, included in our model, have been reported to be highly affected by pH with several species of Gammaproteobacteria particularly abundant in low pH soil [54].

**Figure 4:**
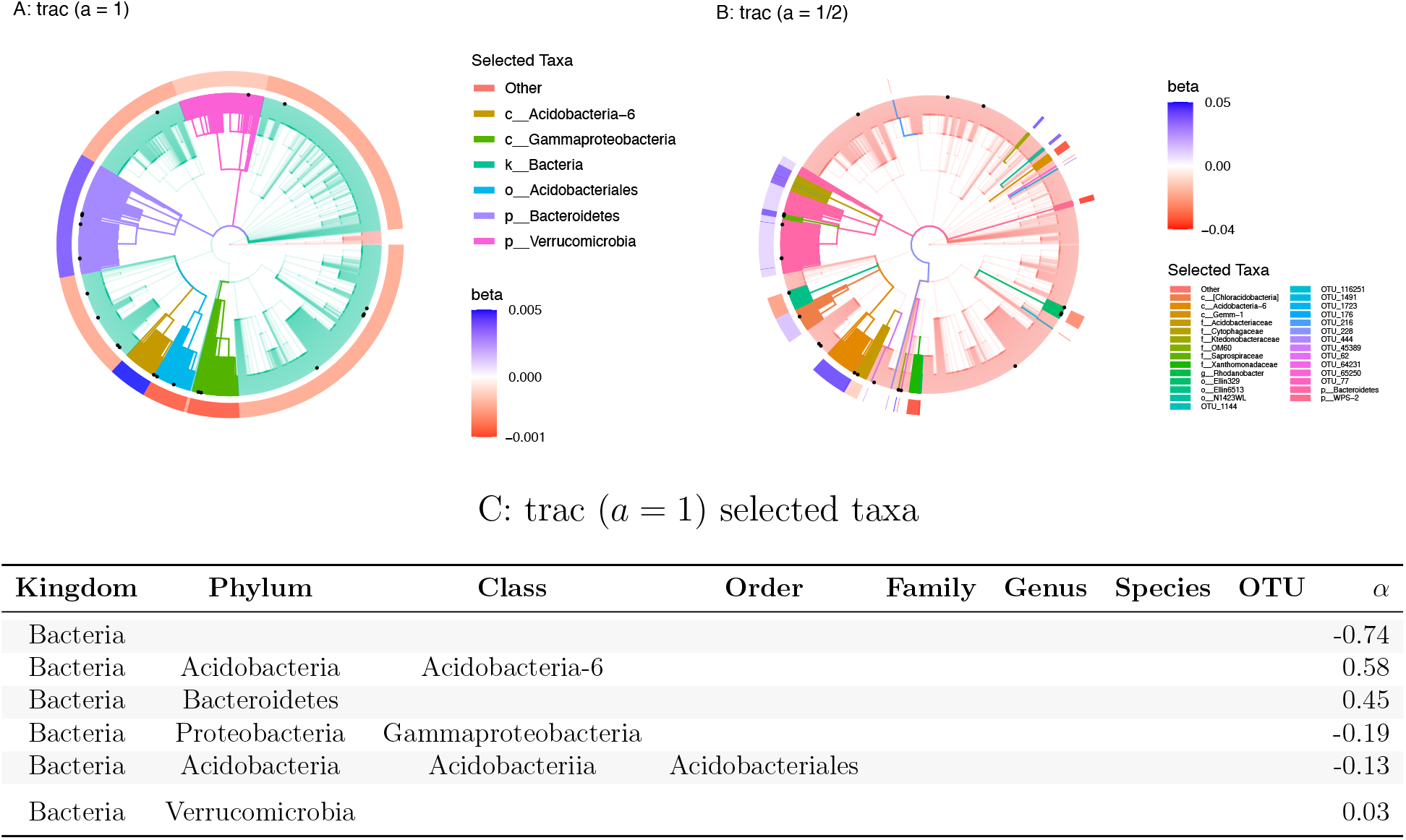
Taxonomic tree visualization of `trac` aggregations (*a* = {1, 1/2} using the Central Park soil data (training/test split 1). Each tree represents the taxonomy of the *p* = 3379 OTUs. Colored branches highlight the estimated `trac` taxon aggregations. The black dots mark the selected taxa of the sparse log-contrast model. The outer rim represents the value of *β* coefficients in the `trac` model from Eq. (1). A: Standard `trac` (*a* = 1) with OTUs as taxon base level selects six aggregations. B: Weighted `trac` (*a* = 1/2) with OTU base level selects 28 aggregations, including 13 on the OTU level. Four of these OTUs are also selected by the sparse log-contrast model which comprises 21 OTUs in total (black dots) (see Suppl. Tables 15 and 16 for the selected coefficients). C: The table lists the *α* coefficients associated with Eq. (2) for the `trac` (*a* = 1) model corresponding to the tree shown in A. These values are in the same units as the pH response variable.

In contrast to the sCD14 data analysis, weighted `trac` (*a* = 1/2) delivers a considerably more fine-grained model with 23 aggregations, including 13 on the OTU level. While the Acidobacteria-6 class is still selected as a whole, weighted `trac` picks specific OTUs and families in the Gammaproteobacteria class. Similar behavior is observed for the Acidobacteriales order and the Bacteroidetes phylum. Moreover, novel orders, families, genera, and OTUs from the Bacteria kingdom are selected. Four OTUs are shared with the sparse log-contrast model which selects 21 OTUs overall.

To compare the models in terms of interpretability and prediction quality, we report in Table 2 average out-of-sample prediction errors and sparsity levels at three different base levels using ten different training/test splits. We observe that for the Central Park soil data set, standard `trac` with OTU base levels delivers the sparsest solutions (on average, ten aggregations), followed by weighted `trac` on the family level (on average, 15 aggregations). The sparse log-contrast models delivers the densest models (26-33, on average). All models are comparable in terms of out-of-sample test error (0.38-0.4).

**Table 2:** Average out-of-sample test errors (rounded average model sparsity in parenthesis) for `trac` (*a* = {1, 1/2}) and sparse log-contrast models, respectively. Each row represents the results for base level OTU, genus, and family. Each value is averaged over ten different training/test splits of the Central Park soil data.

| Base Level | $p$ | **trac** ($a = 1$) | **trac** ($a = 1/2$) | Sparse Log-Contrast |
| --- | --- | --- | --- | --- |
| OTU | 3379 | 0.4 (10) | 0.39 (18) | 0.39 (33) |
| Genus | 2779 | 0.4 (13) | 0.38 (22) | 0.39 (26) |
| Family | 1492 | 0.39 (10) | 0.39 (15) | 0.4 (29) |

### Global predictive model of ocean salinity from Tara data

Integrative marine data collection efforts such as Tara Oceans [55] or the Simons CMAP (https://simonscmap.com) provide the means to investigate ocean ecosystems on a global scale. Using Tara’s environmental and microbial survey of ocean surface water [3], we next illustrate how `trac` can be used to globally connect environmental covariates and marine microbiome data. As an example, we learn global predictive models of ocean salinity from *n* = 136 samples and *p* = 8916 miTAG OTUs [56]. Even though salinity is thought to be an important environmental factor in marine microbial ecosystems, existing studies have investigated the connection between the microbiome and salinity gradients mainly on a local marine scale, in particular estuaries.

Standard `trac` (*a* = 1) identifies four taxonomic aggregations (see Figure 5A), the kingdom Bacteria and the phylum Bacteroidetes being negatively associated and the class Alphaproteobacteria being strongly positively and Gammaproteobacteria being moderately positively associated with marine salinity.

**Figure 5:**
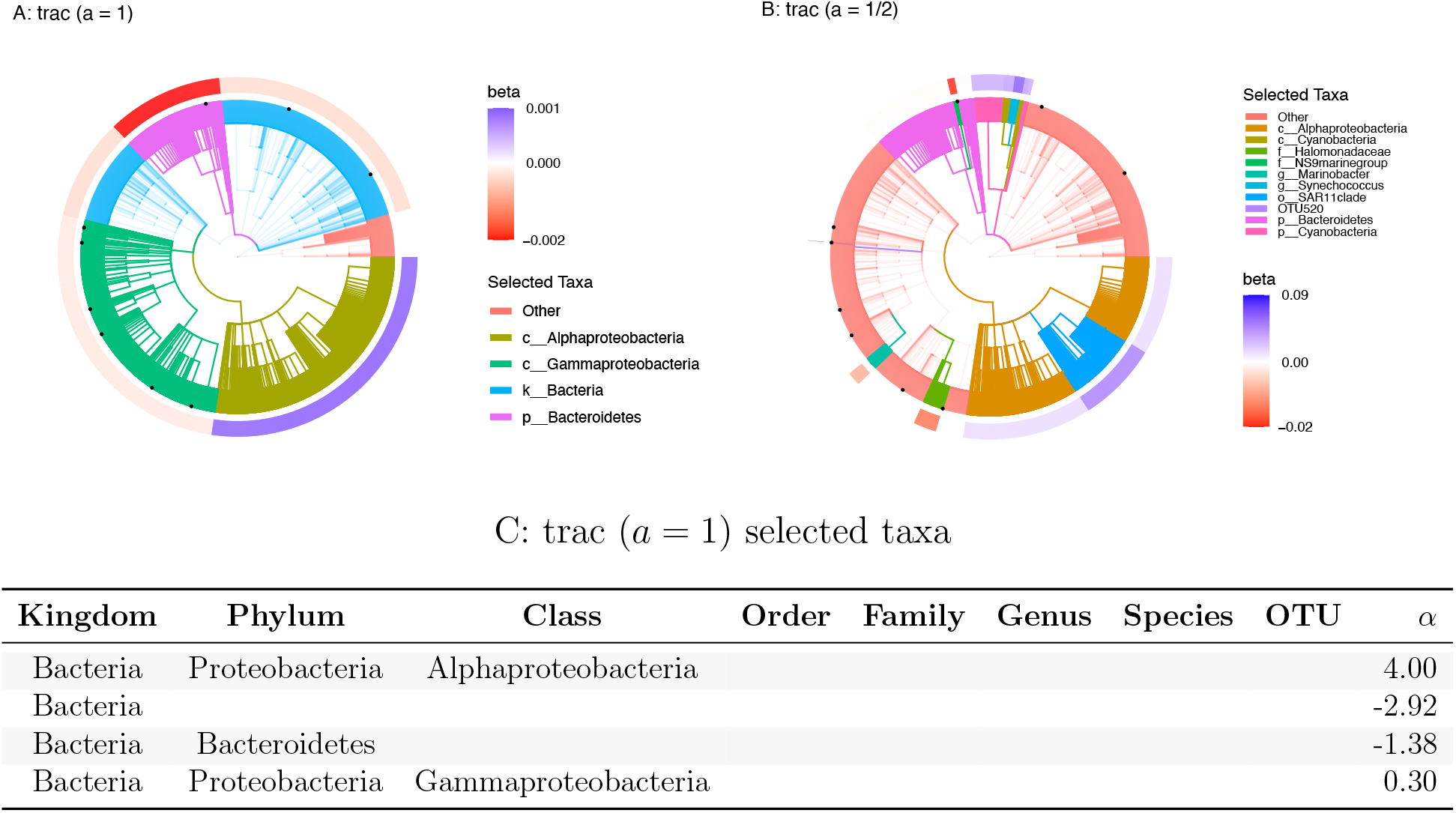
Taxonomic tree visualization of `trac` aggregations (OTUs as taxon base level, *a* = {1, 1/2} for salinity prediction using Tara data (training/test split 1). Each tree represents the taxonomy of the *p* = 8916 miTAG OTUs. Colored branches highlight the estimated `trac` taxon aggregations. The black dots mark the selected taxa of the sparse log-contrast model. The outer rim represents the value of *β* coefficients in the `trac` model from Eq. (1). A: Standard `trac` (*a* = 1) selects four aggregations on the kingdom, phylum, and class level. B: Weighted `trac` (*a* = 1/2) selects ten aggregations across all taxonomic ranks, including a single OTU (OTU520). This OTU is also selected by the sparse log-contrast model which comprises nine OTUs in total (black dots) (see Suppl. Table 18 for the selected coefficients). Both `trac` models select the phylum Bacteroidetes and the Alphaproteobacteria class. C: The table lists the *α* coefficients associated with Eq. (2) for the `trac` (*a* = 1) model corresponding to the tree shown in A. These values are in the same units as the salinity response variable.

Consistent with this `trac` model, a marked increase of Alphaproteobacteria with increasing salinity was observed in several estuary studies [57, 58]. In a global marine microbiome meta-analysis [59], Spearman rank correlations between relative abundances of microbial clades and several physicochemical water properties, including salinity, were reported, showing four out of five orders in the Bacteroidetes phylum to be negatively correlated with salinity. However, three out of four orders belonging to Gammaproteobacteria were negatively correlated with salinity, suggesting that the standard `trac` model does not universally agree with standard univariate assessments. However, as shown in Figure 5B, weighted `trac` (*a* = 1/2) reveals a more fine-grained taxon aggregation, selecting the Halomonadaceae family and the Marinobacter genus in the phylum Gammaproteobacteria to with negative *α* coefficients and a Gammaproteobacteria OTU (OTU 520, order E01-9C-26 marine group) with positive *α* coefficients, respectively (see also Supplementary Table 23). Likewise, out of the nine OTUs selected by the sparse log-contrast model (black dots in Figure 5A,B), four out of six selected Gammaproteobacteria OTUs have negative coefficients (including OTU 520), and two OTUs have positive coefficients.

In terms of model performance, the standard `trac` model shows good global predictive capabilities with an out-of-sample test error of 1.99 (on training/test split 1). We observe, however, that high salinity outliers located in the Red Sea (Coastal Biome) and the Mediterranean Sea (Westerlies Biome) and a low salinity outlier (far eastern Pacific fresh pool south of Panama) are not well captured by the model (see Supplementary Figure 5 for a scatter plot of measured vs. predicted salinity). Weighted `trac` (*a* = 1/2) and the sparse log-contrast models outperform standard `trac` on the salinity prediction task with an out-of-sample test error (on split 1) of 1.94 and 1.52, respectively.

This boost in prediction quality is further confirmed by the average out-of-sample prediction errors across all ten training/test splits and three base levels (see Table 3). Sparse log-contrast models on the OTU and Genus base level perform best (average test error 1.3 and 1.4, respectively), followed by weighted `trac` on Genus level (1.5). However, standard `trac` models are considerably sparser (six to seven aggregations) compared to log-contrast models (13-24 taxa). Weighted `trac` models represent a good trade-off between predictability and interpretability, selecting ten to fourteen taxa, on average.

**Table 3:** Average out-of-sample test errors (rounded average model sparsity in parenthesis) for `trac` (*a* = {1, 1/2}) and sparse log-contrast models, respectively. Each row represents the results for base level OTU, genus, and family and the corresponding dimensionality of the base level. Each value is averaged over ten different training/test splits of the Tara data.

| Base Level | $p$ | **trac** ($a = 1$) | **trac** ($a = 1/2$) | Sparse Log-Contrast |
| --- | --- | --- | --- | --- |
| OTU | 8916 | 2.1 (7) | 1.8 (14) | 1.3 (24) |
| Genus | 4220 | 2 (7) | 1.5 (14) | 1.4 (34) |
| Family | 1869 | 2.1 (6) | 1.7 (10) | 1.6 (13) |

**Table 4:** Average out-of-sample test errors (model sparsity in parenthesis) for `trac` (*a* = {1, 1/2}) and sparse log-contrast models, respectively. Each row considers a different base level (OTU, genus, and family). Each number is averaged over ten different training/test splits of the Fram Strait (FL) data.

| Base Level | $p$ | **trac** ($a = 1$) | **trac** ($a = 1/2$) | Sparse Log-Contrast |
| --- | --- | --- | --- | --- |
| OTU | 4510 | 1.9e+02 (2) | 1.5e+02 (5) | 1.7e+02 (4) |
| Genus | 2930 | 1.9e+02 (3) | 1.5e+02 (4) | 1.4e+02 (6) |
| Family | 1125 | 1.8e+02 (4) | 1.4e+02 (4) | 1.5e+02 (4) |

**Table 5:** Coefficients selected by `trac` (a = 1) for Gut (HIV): sCD14

| Kingdom | Phylum | Class | Order | Family | Genus | Species | OTU | $\alpha$ |
| --- | --- | --- | --- | --- | --- | --- | --- | --- |
| Bacteria | Firmicutes | Clostridia | Clostridiales | Ruminococcaceae |  |  |  | 2221.75 |
| Bacteria | Firmicutes | Clostridia | Clostridiales | Lachnospiraceae |  |  |  | -1644.86 |
| Bacteria | Actinobacteria |  |  |  |  |  |  | -501.43 |
| Bacteria |  |  |  |  |  |  |  | -362.27 |
| Bacteria | Bacteroidetes | Bacteroidia | Bacteroidales | Bacteroidaceae | Bacteroides |  |  | 286.80 |

**Table 6:** Coefficients selected by `trac` (a = 1/2) for Gut (HIV): sCD14

| Kingdom | Phylum | Class | Order | Family | Genus | Species | OTU | $\alpha$ |
| --- | --- | --- | --- | --- | --- | --- | --- | --- |
| Bacteria | Firmicutes | Clostridia | Clostridiales | Ruminococcaceae |  |  |  | 629.10 |
| Bacteria | Actinobacteria |  |  |  |  |  |  | -570.60 |
| Bacteria | Firmicutes | Negativicutes | Selenomonadales | Veillonellaceae | Mitsuokella | - | Otu000070 | -128.83 |
| Bacteria | Firmicutes | Clostridia | Clostridiales | Lachnospiraceae | Lachnospira |  |  | -125.49 |
| Bacteria | Firmicutes | Clostridia | Clostridiales | Ruminococcaceae | Subdoligranulum |  |  | 121.80 |
| Bacteria | Bacteroidetes | Bacteroidia | Bacteroidales | Bacteroidaceae | Bacteroides |  |  | 82.62 |
| Bacteria | Bacteroidetes | Bacteroidia | Bacteroidales | Bacteroidaceae | Bacteroides | - | Otu000014 | 51.81 |
| Bacteria | Firmicutes | Clostridia | Clostridiales | Lachnospiraceae | Lachnospira | - | Otu000038 | -49.69 |
| Bacteria | Bacteroidetes | Bacteroidia | Bacteroidales | Prevotellaceae | Alloprevotella | - | Otu000011 | 41.31 |
| Bacteria | Firmicutes | Clostridia | Clostridiales | Ruminococcaceae | Incertae_Sedis | - | Otu000073 | -39.42 |
| Bacteria | Actinobacteria | Actinobacteria | Bifidobacteriales | Bifidobacteriaceae | Bifidobacterium | - | Otu000098 | -12.61 |

**Table 7:**
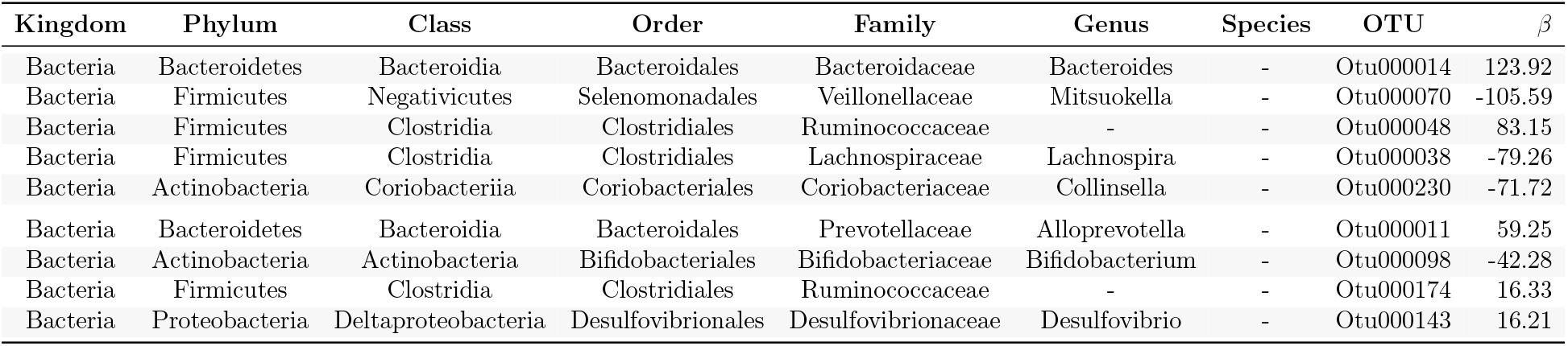
Coefficients selected by the sparse log-contrast method for Gut (HIV): sCD14

**Table 8:**
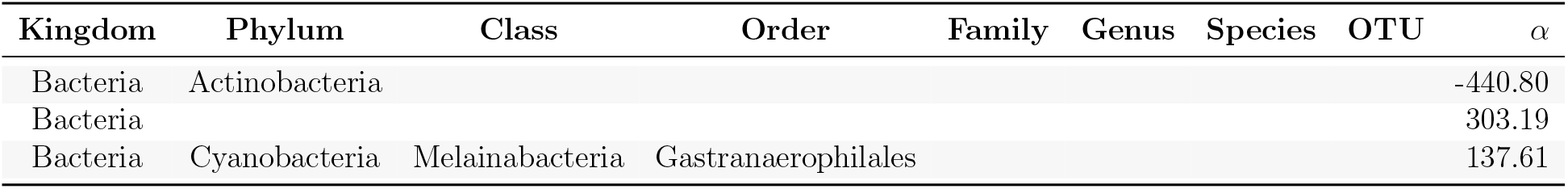
Coefficients selected by `trac` on family level (a = 1) for Gut (HIV): sCD14

| Kingdom | Phylum | Class | Order | Family | Genus | Species | OTU | $\alpha$ |
| --- | --- | --- | --- | --- | --- | --- | --- | --- |
| Bacteria | Actinobacteria |  |  |  |  |  |  | -440.80 |
| Bacteria |  |  |  |  |  |  |  | 303.19 |
| Bacteria | Cyanobacteria | Melainabacteria | Gastranaerophilales |  |  |  |  | 137.61 |

**Table 9:** Coefficients selected by `trac` on family level (a = 1/2) for Gut (HIV): sCD14

| Kingdom | Phylum | Class | Order | Family | Genus | Species | OTU | $\alpha$ |
| --- | --- | --- | --- | --- | --- | --- | --- | --- |
| Bacteria | Actinobacteria |  |  |  |  |  |  | -419.10 |
| Bacteria |  |  |  |  |  |  |  | 301.98 |
| Bacteria | Proteobacteria | Gammaproteobacteria | Enterobacteriales | Enterobacteriaceae |  |  |  | 112.87 |
| Bacteria | Bacteroidetes | Bacteroidia | Bacteroidales | Bacteroidaceae |  |  |  | 9.42 |
| Bacteria | Proteobacteria | Gammaproteobacteria | Aeromonadales | Succinivibrionaceae |  |  |  | -5.18 |

**Table 10:** Coefficients selected by the sparse log-contrast method on family level for Gut (HIV): sCD14

| Kingdom | Phylum | Class | Order | Family | Genus | Species | OTU | $\beta$ |
| --- | --- | --- | --- | --- | --- | --- | --- | --- |
| Life | Bacteria | Actinobacteria | Coriobacteriia | Coriobacteriales | Coriobacteriaceae |  |  | -317.40 |
| Life | Bacteria | Proteobacteria | Gammaproteobacteria | Enterobacteriales | Enterobacteriaceae |  |  | 177.47 |
| Life | Bacteria | Cyanobacteria | Melainabacteria | Gastranaerophilales | [Unclassified] |  |  | 138.24 |
| Life | Bacteria | Actinobacteria | Actinobacteria | Bifidobacteriales | Bifidobacteriaceae |  |  | -68.26 |
| Life | Bacteria | Firmicutes | Clostridia | Clostridiales | Defluviitaleaceae |  |  | 44.07 |
| Life | Bacteria | Proteobacteria | Alphaproteobacteria | [Unclassified] | [Unclassified] |  |  | 25.88 |

## Conclusions

Finding predictive and interpretable relationships between microbial amplicon sequencing data and ecological, environmental, or host-associated covariates of interest is a cornerstone of exploratory data analysis in microbial biogeography and ecology. To this end, we have introduced `trac`, a scalable tree-aggregation regression framework for compositional amplicon data. The framework leverages the hierarchical nature of microbial sequencing data to learn parsimonious log-ratios of microbial compositions along the taxonomic or phylogenetic tree that best predict continuous environmental or host-associated response variables. The `trac` method is applicable to any user-defined taxon base level as input, e.g., ASV/OTU, genus, or family level, and includes a scalar tuning parameter a that allows control of the overall aggregation granularity. As shown above, this allows seamless testing of a continuum of models to a data set of interest, with prior approaches to sparse log-contrast modeling modeling as special limit cases [15, 60, 43]. The framework, available in the R package `trac` and Python [35], shares similarities with ideas from tree-guided, *balance* modeling of compositional data [18, 24, 23], albeit with a stronger focus on finding *predictive* relationships and emphasis on fast computation thanks to the convexity of the formulation and the underlying efficient path algorithm.

Our comprehensive benchmarks and comparative analysis on host-associated and environmental microbiome data revealed several notable observations. Firstly, across almost all tested taxon base levels and methods, standard `trac` (*a* = 1) resulted in the most parsimonious models and revealed data-specific taxon aggregations comprising all taxonomic orders. This facilitated straightforward model interpretability despite the high-dimensionality of the data. For instance, on the sCD14 data, the standard `trac` model with OTU base level asserted a particularly strong predictive role of the Ruminococcaceae/Lachnospiraceae family ratio for sCD14, thus generating testable biological hypothesis. Likewise, `trac` analysis on environmental microbiomes in soil and marine habitats consistently provided parsimonious taxonomic aggregations for predicting covariates of interest. For instance, Alpha- and Gammaproteobacteria/Bacteroidetes ratios well-aligned with sea surface water salinity on a global scale, reminiscent of the ubiquitous Firmicutes/Bacteroidetes ratio in the context of the gut microbiome and obesity [61, 62].

Secondly, arithmetic aggregation of OTUs to a higher taxonomic base level prior to `trac` or sparse log-contrast modeling did not result in significant predictive performance gains. In fact, using OTUs as base level, at least one of the three statistical methods showed superior test error performance while maintaining a high level of sparsity. These results suggest that a user may safely choose the highest level of resolution of the data (e.g., mOTUs, OTUs, or ASVs) in (weighted) `trac` models without sacrificing prediction performance.

Thirdly, while standard `trac` models always showed good predictive performance on out-of-sample test data, our comparative and average analysis indicated that weighted `trac` and sparse log-contrast models can outperform the parsimonious `trac` models in terms of test error, particularly on environmental microbiome data. For instance, on Central Park soil data, we observed moderate performance gains using weighted `trac`, and on marine data (see Extended Results in the Supplementary Material for the Fram Strait dataset), sparse log-contrast models showed, on average, the best predictive performance. These results add a valuable piece of information to the ongoing debate about the usefulness of incorporating phylogenetic or taxonomic information into statistical modeling. For example, the authors in [63] convincingly argue that incorporating such information provides no gains in microbial differential abundance testing scenarios.

We posit that, in the context of statistical regression, full comparative `trac` analyses like the ones presented here, can immediately determine in a concrete and objective way whether phylogenetic or taxonomic information is useful for a particular prediction task on the data set of interest.

The `trac` framework naturally lends itself to several methodological extensions that are easy to implement and may prove valuable in microbiome research. Firstly, as apparent in the gut microbiome context, inclusion of additional factors such as diet and life style would likely improve prediction performance. This can be addressed by combining `trac` with standard (sparse) linear regression to allow the incorporation of (non-compositional) covariates into the statistical model (see, e.g., [64]). Secondly, while we focused on predictive regression modeling of continuous outcomes, it is straightforward to adopt our framework to classification tasks when binary outcomes, such as, e.g., case vs. control group, or healthy vs. sick participants, are to be predicted. For instance, using the (Huberized) square hinge loss (see, e.g., [65]) as objective function *L*(·) in Eq. (2) would provide an ideal means to handle binary responses while simultaneously enabling the use of efficient path algorithms (see [35] and references therein). Thirdly, due to the compositional nature of current amplicon data, we presented `trac` in the common framework of log-contrast modeling. However, alternative forms of tree aggregations over compositions are possible, for instance, by directly using the relative abundances as features rather than log-transformed quantities. Tree aggregations would then amount to grouped relative abundance *differences* and not log-ratios, thus resulting in a different interpretation of the estimated model features.

In summary, we believe that our methodology and its implementation in the R package `trac`, together with the presented reproducible application workflows, provide a valuable blueprint for future data-adaptive aggregation and regression modeling for microbial biomarker discovery, biogeography, and ecology research. This, in turn, may contribute to the generation of new interpretable and testable hypotheses about host-microbiome interactions and the general factors that shape microbial ecosystems in their natural habitats.

## Acknowledgments

We thank Dr. Javier Rivera-Pinto for providing the processed OTU tables for the Gut (HIV)-sCD14 data set. This work was supported by the Simons Collaboration on Computational Biogeochemical Modeling of Marine Ecosystems/CBIOMES (Grant ID: 549939, JB). Jacob Bien was also supported in part by NIH Grant R01GM123993 and NSF CAREER Award DMS-1653017. Christian L. Müller acknowledges support from the Center for Computational Mathematics, Flatiron Institute, and the Institute of Computational Biology, Helmholtz Zentrum München. An earlier version of this work appeared as Chapter 4 of Xiaohan Yan’s PhD dissertation [66].

## Supplementary Material

### A Data and Code availability

The data and code for fully reproducing all results presented in this manuscript are available at Zenodo at https://doi.org/10.5281/zenodo.4734527. The simulation code has been tested on R version 4.0. The `trac` R package is available at https://github.com/jacobbien/trac. A vignette describing key functionalities of the package and an archetypical workflow are available at https://jacobbien.github.io/trac/articles/trac-example.html. The `c-lasso` Python package [1] is available at https://github.com/Leo-Simpson/c-lasso and can be installed via `pip`.

### B Derivation of Optimization Problem

We design a convex tree-based penalty 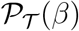 that promotes *β* to be constant along branches of 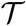. We encode 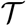 through a binary matrix 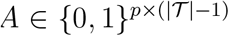 indicating whether feature *j* is a leaf of each non-root node 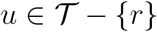, that is 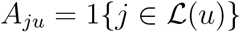 where 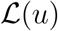 is the set of leaves that descend from *u*. In particular, we take

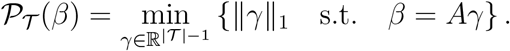

**Figure 1:**
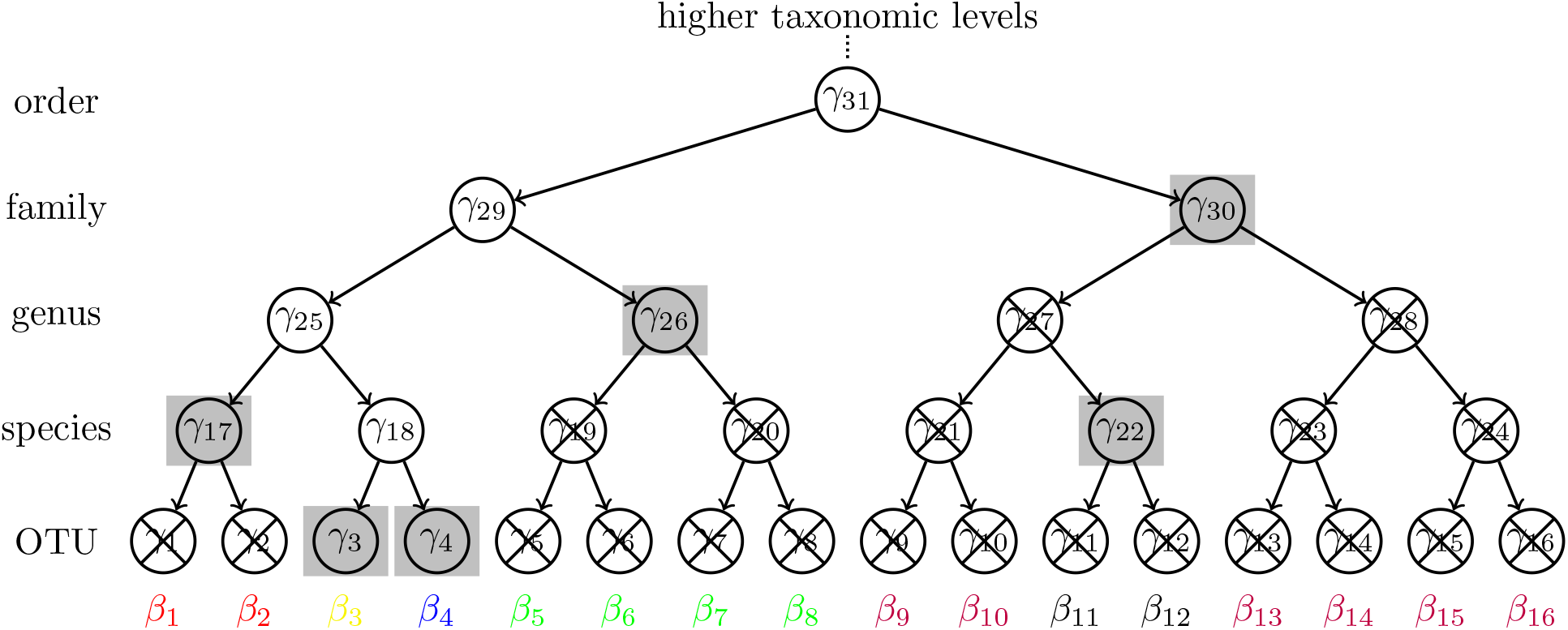
Schematic of the tree aggregation process.

Figure 1 shows a schematic of the tree aggregation idea. The vector 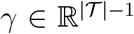 can be thought of as a latent parameter vector with an entry associated with each node of the tree (see Figure 1). We associate a *β_j_* to each leaf of 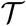, and the constraint *β* = *Aγ* expresses a particular relationship between these, namely that each coefficient *β_j_* is the sum of the *γ_u_* for which 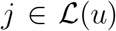 (i.e., each *β_j_* is the sum of its ancestor *γ*-values in the tree). This relationship implies that when all the *γ*-values in a subtree are zero (denoted by crossed out nodes in the figure), then all the *β* coefficients within the subtree are equal. Thus, the sparsity inducing ℓ_1_-norm on γ in 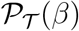 induces *β* to tend to be constant within subtrees of 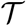. Using this penalty in Eq. (1) in the main paper leads to the `trac` method, which is computed by solving,

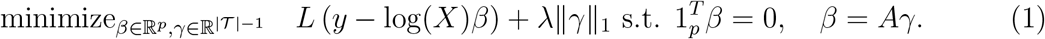

This estimator is built on the tree-based aggregation penalty in [2], developed for general situations in which features are rare and a tree relating the features is available. In their setting, features are not compositional, so they do not introduce a sum-to-zero constraint or take the log of the features. The `trac` problem can be written more simply, entirely in terms of *γ*, as

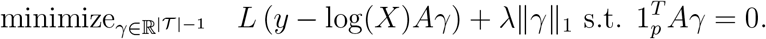

The 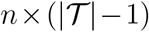 matrix log(*X*)*a* has the sum of the log counts of each of the 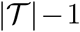 subtrees of 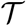 (excluding 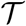 itself). Changing variables to 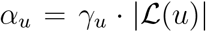 and using properties of logarithms establishes the equivalence with problem Eq. (2) in the main paper.

### C Extended Results

We provide extended results, including an in-depth analysis of `trac` prediction of BMI from American Gut Project data, moisture prediction in Central Park soil, and leucine prediction in the Fram Strait.

#### Immune marker sCD14 prediction in HIV patients

For the sCD14 data, we provide coefficient tables learned by `trac` (*a* = 1), `trac` (*a* = 1/2), and the sparse log-contrast model on the first random train-test data split (of ten) in Section D. This complements the tree visualizations shown in the main manuscript. We also include the results on the family base level (corresponding to panels C and D of Figure 3 in the main paper).

#### BMI prediction from American Gut microbiome profiles

Finding consistent gut microbial signatures that are predictive of a person’s body mass index (BMI) remains a non-trivial problem. Several early studies argued that obesity is associated with phylum-level changes in the microbiome [3], including increased Firmicutes to Bacteroidetes phyla ratios [4], often referred to as a hallmark predictor of obesity. The authors in [5] and [6] were among the first to identify a small set of microbial genera that were (moderately) predictive of host BMI using sparse log-contrast models on the COMBO microbiome dataset [7].

Using `trac`, we revisit BMI prediction from microbial abundance data using a subset of the American Gut Project (AGP) data comprising *p* = 1387 OTUs across *n* = 6266 participants in the lean to obese BMI range. The standard `trac` model (*a* =1) with the 1SE rule identified a model with 132 predictors, consisting of aggregations across *all* taxonomic levels. Table 11 summarizes the 15 strongest predictors which include the kingdom Bacteria (vs. Archaea) as negative baseline, the phylum Bacteroidetes and several families and genera in the class Clostridia (which belongs to the Firmicutes phylum) with positive associations. The strongest positive OTU level predictor is an unknown species belonging to the Ruminococcaceae family. Figure 2 shows the corresponding `trac` model BMI predictions (with 1SE rule) vs. measured BMI on the test set (split 1). The out-of-sample test error on this split is 15.31, and roughly 16 on average across all ten splits (see Table 1). Standard `trac`, weighted `trac`, and sparse log-contrast models show similar performance in terms of test error (16 – 17) across all taxon base levels, with sparsity levels between 73 and 122 on OTU and genus level, and about 23-27 on the family level.

**Table 11:** Top 15 coefficients selected by `trac` (a = 1) for Gut (AGP): BMI

| Kingdom | Phylum | Class | Order | Family | Genus | Species | OTU | $\alpha$ |
| --- | --- | --- | --- | --- | --- | --- | --- | --- |
| Bacteria |  |  |  |  |  |  |  | -11.95 |
| Bacteria | Firmicutes | Clostridia | Clostridiales | Ruminococcaceae |  |  |  | 2.86 |
| Bacteria | Firmicutes | Clostridia | Clostridiales | Lachnospiraceae |  |  |  | 2.23 |
| Bacteria | Bacteroidetes |  |  |  |  |  |  | 1.45 |
| Bacteria | Firmicutes | Clostridia | Clostridiales |  |  |  |  | 1.18 |
| Bacteria | Proteobacteria | Gammaproteobacteria | Enterobacteriales | Enterobacteriaceae |  |  |  | 0.90 |
| Bacteria | Firmicutes | Erysipelotrichi | Erysipelotrichales | Erysipelotrichaceae |  |  |  | -0.80 |
| Bacteria | Firmicutes | Clostridia | Clostridiales | Lachnospiraceae | Blautia |  |  | 0.73 |
| Bacteria | Firmicutes | Bacilli | Lactobacillales |  |  |  |  | 0.72 |
| Bacteria | Firmicutes | Clostridia | Clostridiales | Veillonellaceae |  |  |  | 0.71 |
| Bacteria | Firmicutes | Clostridia | Clostridiales | Lachnospiraceae | Dorea |  |  | 0.51 |
| Bacteria | Firmicutes | Clostridia | Clostridiales | Ruminococcaceae | Ruminococcus |  |  | 0.49 |
| Bacteria | Firmicutes | Clostridia | Clostridiales | [Mogibacteriaceae] |  |  |  | -0.36 |
| Bacteria | Bacteroidetes | Bacteroidia | Bacteroidales | [Barnesiellaceae] |  |  |  | 0.32 |
| Bacteria | Firmicutes | Clostridia | Clostridiales | Ruminococcaceae | - | - | 4356062 | 0.30 |

**Table 12:** Top 15 coefficients selected by `trac` (a = 1/2) for Gut (AGP): BMI

| Kingdom | Phylum | Class | Order | Family | Genus | Species | OTU | $\alpha$ |
| --- | --- | --- | --- | --- | --- | --- | --- | --- |
| Bacteria | Firmicutes | Erysipelotrichi | Erysipelotrichales | Erysipelotrichaceae |  |  |  | -0.30 |
| Bacteria | Firmicutes | Clostridia | Clostridiales | Ruminococcaceae | - | - | 4356062 | 0.28 |
| Bacteria | Proteobacteria | Gammaproteobacteria | Enterobacteriales | Enterobacteriaceae |  |  |  | -0.24 |
| Bacteria | Firmicutes | Bacilli | Lactobacillales |  |  |  |  | 0.23 |
| Bacteria | Proteobacteria | Gammaproteobacteria | Pasteurellales | Pasteurellaceae | Haemophilus | parainfluenzae | 4477696 | -0.21 |
| Bacteria | Firmicutes | Clostridia | Clostridiales | Lachnospiraceae | Dorea | - | 181871 | 0.19 |
| Bacteria | Firmicutes | Clostridia | Clostridiales | Lachnospiraceae | Blautia | - | 4361189 | 0.19 |
| Bacteria | Firmicutes | Clostridia | Clostridiales | [Tissierellaceae] | Finegoldia | - | 1096610 | 0.17 |
| Bacteria | Firmicutes | Erysipelotrichi | Erysipelotrichales | Erysipelotrichaceae | Catenibacterium | - | 4480861 | 0.16 |
| Bacteria | Actinobacteria | Actinobacteria |  |  |  |  |  | -0.15 |
| Bacteria | Firmicutes | Erysipelotrichi | Erysipelotrichales | Erysipelotrichaceae | - | - | 145801 | -0.14 |
| Bacteria | Firmicutes | Clostridia | Clostridiales | - | - | - | 195004 | 0.14 |
| Bacteria | Bacteroidetes | Bacteroidia | Bacteroidales | [Barnesiellaceae] |  |  |  | 0.13 |
| Bacteria | Firmicutes | Bacilli | Bacillales | Staphylococcaceae | Staphylococcus | aureus | 4446058 | -0.13 |
| Bacteria | Actinobacteria | Coriobacteriia | Coriobacteriales | Coriobacteriaceae | Eggerthella | lenta | 4393532 | -0.12 |

**Table 13:**
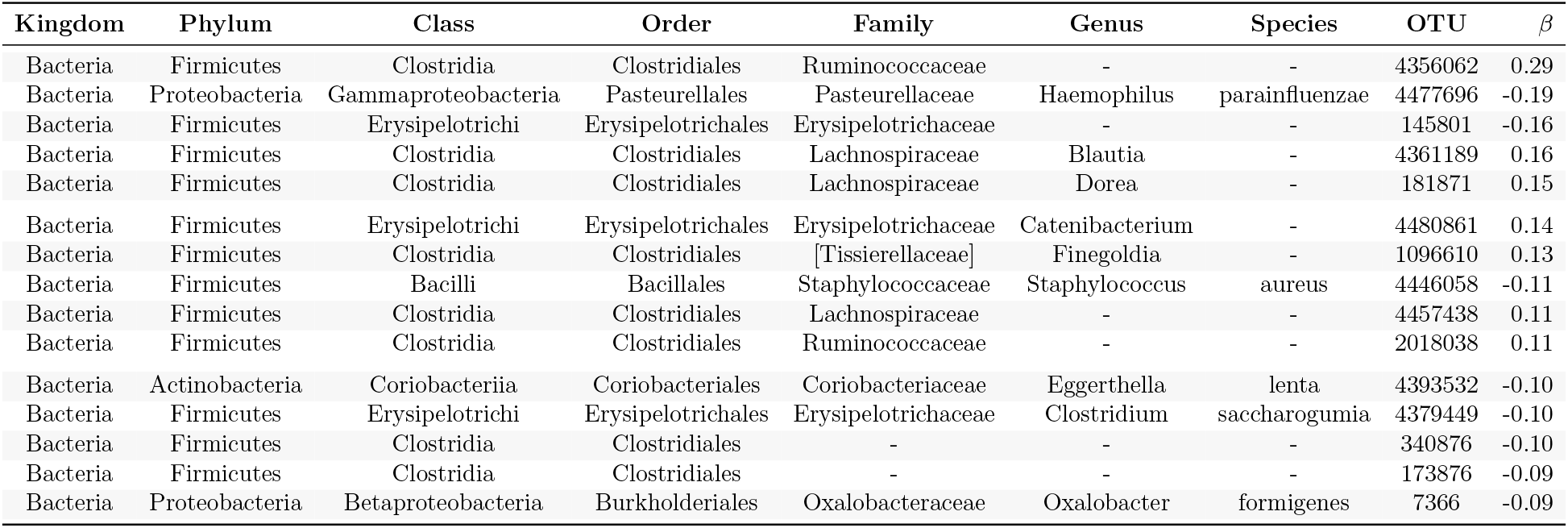
Top 15 coefficients selected by the sparse log-contrast method for Gut (AGP): BMI

| Kingdom | Phylum | Class | Order | Family | Genus | Species | OTU | $\beta$ |
| --- | --- | --- | --- | --- | --- | --- | --- | --- |
| Bacteria | Firmicutes | Clostridia | Clostridiales | Ruminococcaceae | - | - | 4356062 | 0.29 |
| Bacteria | Proteobacteria | Gammaproteobacteria | Pasteurellales | Pasteurellaceae | Haemophilus | parainfluenzae | 4477696 | -0.19 |
| Bacteria | Firmicutes | Erysipelotrichi | Erysipelotrichales | Erysipelotrichaceae | - | - | 145801 | -0.16 |
| Bacteria | Firmicutes | Clostridia | Clostridiales | Lachnospiraceae | Blautia | - | 4361189 | 0.16 |
| Bacteria | Firmicutes | Clostridia | Clostridiales | Lachnospiraceae | Dorea | - | 181871 | 0.15 |
| Bacteria | Firmicutes | Erysipelotrichi | Erysipelotrichales | Erysipelotrichaceae | Catenibacterium | - | 4480861 | 0.14 |
| Bacteria | Firmicutes | Clostridia | Clostridiales | [Tissierellaceae] | Finegoldia | - | 1096610 | 0.13 |
| Bacteria | Firmicutes | Bacilli | Bacillales | Staphylococcaceae | Staphylococcus | aureus | 4446058 | -0.11 |
| Bacteria | Firmicutes | Clostridia | Clostridiales | Lachnospiraceae | - | - | 4457438 | 0.11 |
| Bacteria | Firmicutes | Clostridia | Clostridiales | Ruminococcaceae | - | - | 2018038 | 0.11 |
| Bacteria | Actinobacteria | Coriobacteriia | Coriobacteriales | Coriobacteriaceae | Eggerthella | lenta | 4393532 | -0.10 |
| Bacteria | Firmicutes | Erysipelotrichi | Erysipelotrichales | Erysipelotrichaceae | Clostridium | saccharogumia | 4379449 | -0.10 |
| Bacteria | Firmicutes | Clostridia | Clostridiales | - | - | - | 340876 | -0.10 |
| Bacteria | Firmicutes | Clostridia | Clostridiales | - | - | - | 173876 | -0.09 |
| Bacteria | Proteobacteria | Betaproteobacteria | Burkholderiales | Oxalobacteraceae | Oxalobacter | formigenes | 7366 | -0.09 |

**Table 14:** Coefficients selected by `trac` (a = 1) for Central Park Soil: pH

| Kingdom | Phylum | Class | Order | Family | Genus | Species | OTU | $\alpha$ |
| --- | --- | --- | --- | --- | --- | --- | --- | --- |
| Bacteria |  |  |  |  |  |  |  | -0.74 |
| Bacteria | Acidobacteria | Acidobacteria-6 |  |  |  |  |  | 0.58 |
| Bacteria | Bacteroidetes |  |  |  |  |  |  | 0.45 |
| Bacteria | Proteobacteria | Gammaproteobacteria |  |  |  |  |  | -0.19 |
| Bacteria | Acidobacteria | Acidobacteriia | Acidobacteriales |  |  |  |  | -0.13 |
| Bacteria | Verrucomicrobia |  |  |  |  |  |  | 0.03 |

The standard `trac` model contains aggregations across all taxonomic levels. For instance, on the genus level, `trac` selects Blautia, Dorea, and Ruminococcus as positive predictors. The strongest overall positive predictors are the Bacteroidetes phylum, and the Ruminococcaceae, Lachnospiraceae, and Clostridiales families. The Lachnospiraceae/Bacteria ratio is also the first log-contrast to enter the `trac` aggregation path on the AGP data. The Erysipelotrichaceae and the Mogibacteriaceae families are the strongest negative predictors. Consistent with our model, Mogibacteriaceae were shown to be more abundant in lean individuals [8], and Erysipelotrichaceae were recently reported to be more abundant in normal compared to obese people or subjects with metabolic disorder [9]. However, the fact that standard `trac` could not identify a simple sparse predictive aggregation model for BMI suggests that more complex statistical models are required for predictive modeling, including adjustment for available covariates such as diet, sex, and overall life style.

**Figure 2:**
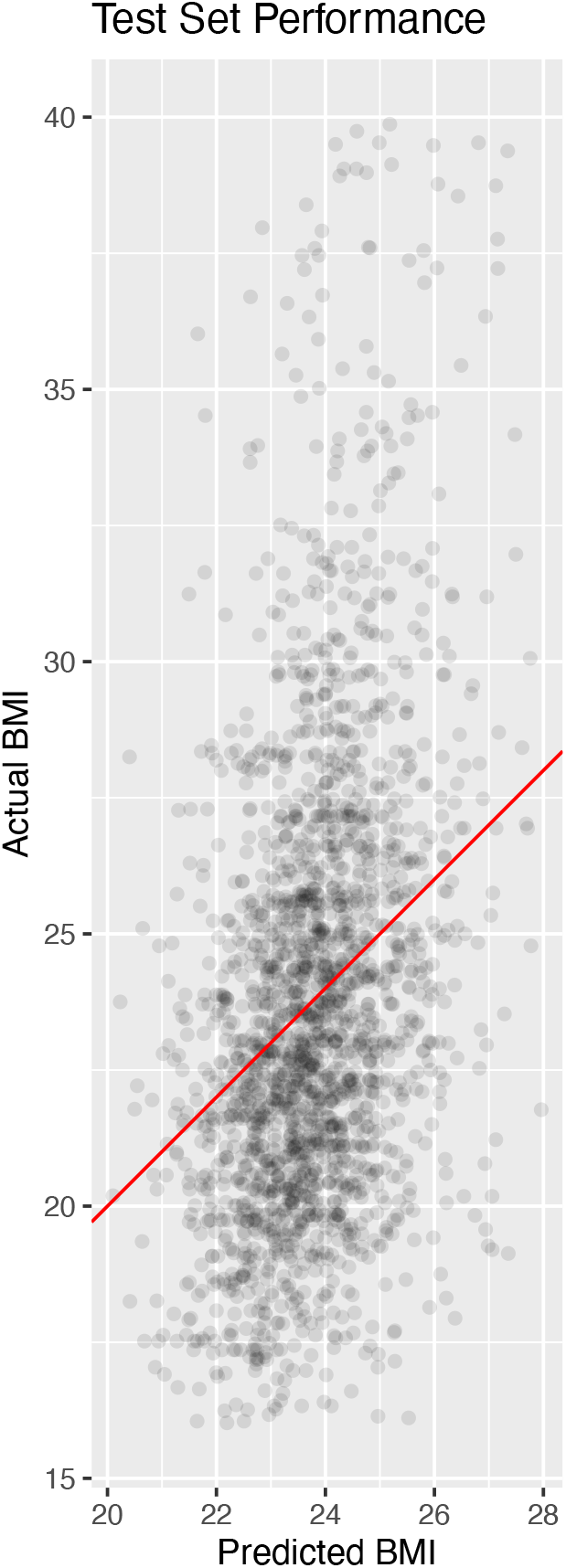
A scatter plot of measured BMI (y-axis) vs. `trac` model BMI predictions on a test set of *n* = 2088 AGP participants shows that predicted BMIs largely cover the “normal” BMI range between 20 and 28 with an overall test set correlation of 0.33. This model has 132 selected taxa, ranging from Kingdom to OTU levels. Table 11 shows the top 15 aggregations with largest *α*-coefficients.

**Table 1:** Average out-of-sample test errors (model sparsity in parenthesis) for `trac` (*a* = {1, 1/2}) and sparse log-contrast models, respectively. Each row considers a different base level (OTU, genus, and family). Each number is averaged over ten different training/test splits of the Gut (AGP), BMI data.

| Base Level | $p$ | **trac** ($a = 1$) | **trac** ($a = 1/2$) | Sparse Log-Contrast |
| --- | --- | --- | --- | --- |
| OTU | 1387 | 16 (115) | 16 (100) | 16 (81) |
| Genus | 824 | 16 (73) | 16 (111) | 16 (122) |
| Family | 199 | 17 (27) | 17 (23) | 17 (24) |

#### Predicting Central Park pH and soil moisture from microbial communities

Here, we complement the microbiome-pH analysis from the main text with an investigation of the relationship between soil microbiome and gravimetric moisture (% water) measurements in Central Park. Since pH and moisture measurements are uncorrelated in the Central Park dataset, we also investigated the similarity between the predictive aggregations for pH and moisture.

Standard `trac` inferred a predictive model of moisture consisting of 23 taxonomic aggregations, including the phylum Proteobacteria and the classes Alpha- and Deltaproteobacteria as strong positive predictors, and the phyla Verrucomicrobia, Actinobacteria, and the order Sphingobacteriales as strong negative predictors (see Table 15). On the test data (split 1), the correlation between model predictions and measurements was 0.42. Compared to pH, the reduced predictive power is in agreement with [10]’s observation about the smaller influence of SMD compared to pH on microbial composition. Nonetheless, `trac`’s taxonomic groupings provide meaningful information about the taxonomic structure of soil microbiota along moisture gradients. For example, the model supports the positive association between Proteobacteria and moisture, as previously observed in a study along a vegetation gradient on the Loess Plateau in China [11], and the negative effect of moisture on the phylum Verrucomicrobia and the positive effect on Deltaproteobacteria in the Giessen free-air CO2 enrichment (Gi-FACE) experiment [12]. The Gi-FACE study, however, also reported several relationships between the microbiome and the soil moisture that are incongruent with our model, including the role of Acidobacteria.

**Figure 3:**
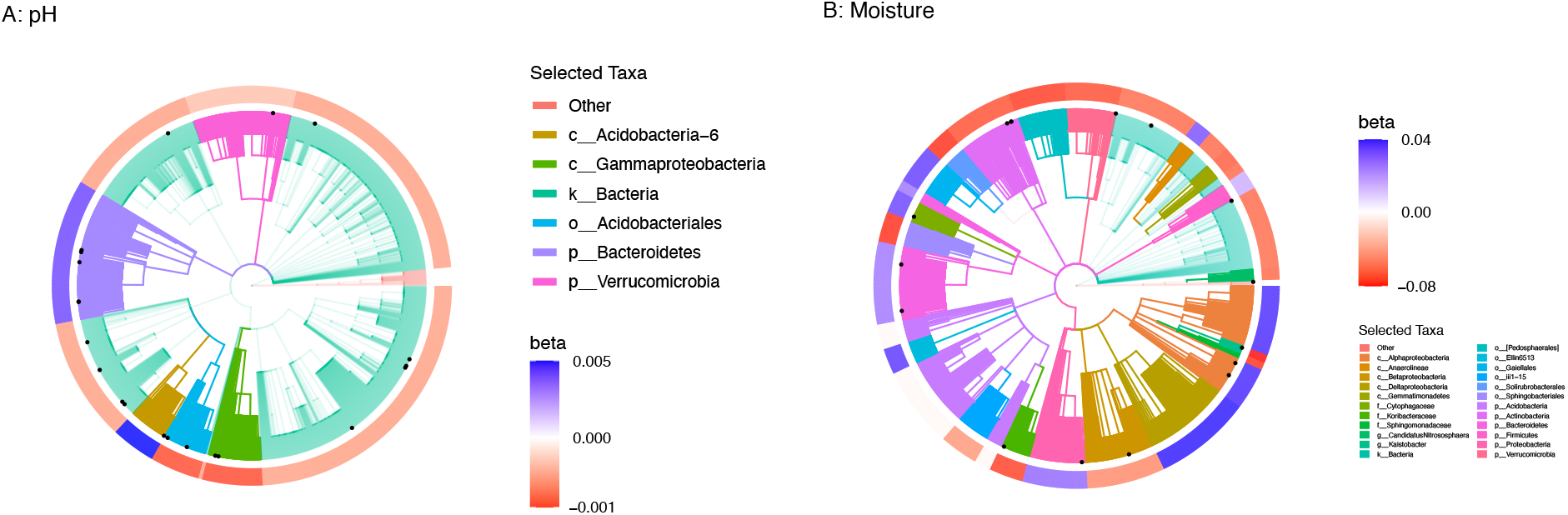
Taxonomic aggregations (as highlighted by branch colors) inferred by `trac` (*a* = 1), that are predictive of Central Park soil pH and moisture, respectively. The color coding on the outermost ring corresponds to the estimated leaf coefficients *β* and are in units of the response (which differs in the two cases).

**Table 15:**
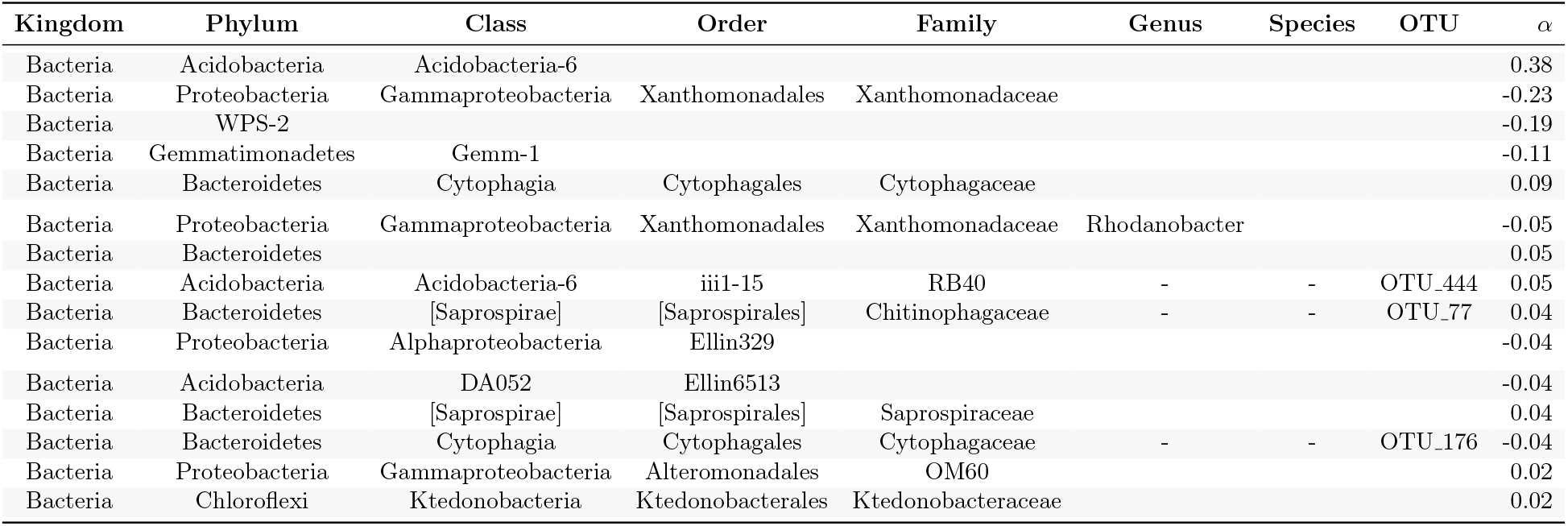
Top 15 coefficients selected by `trac` (a = 1/2) for Central Park Soil: pH

**Table 16:**
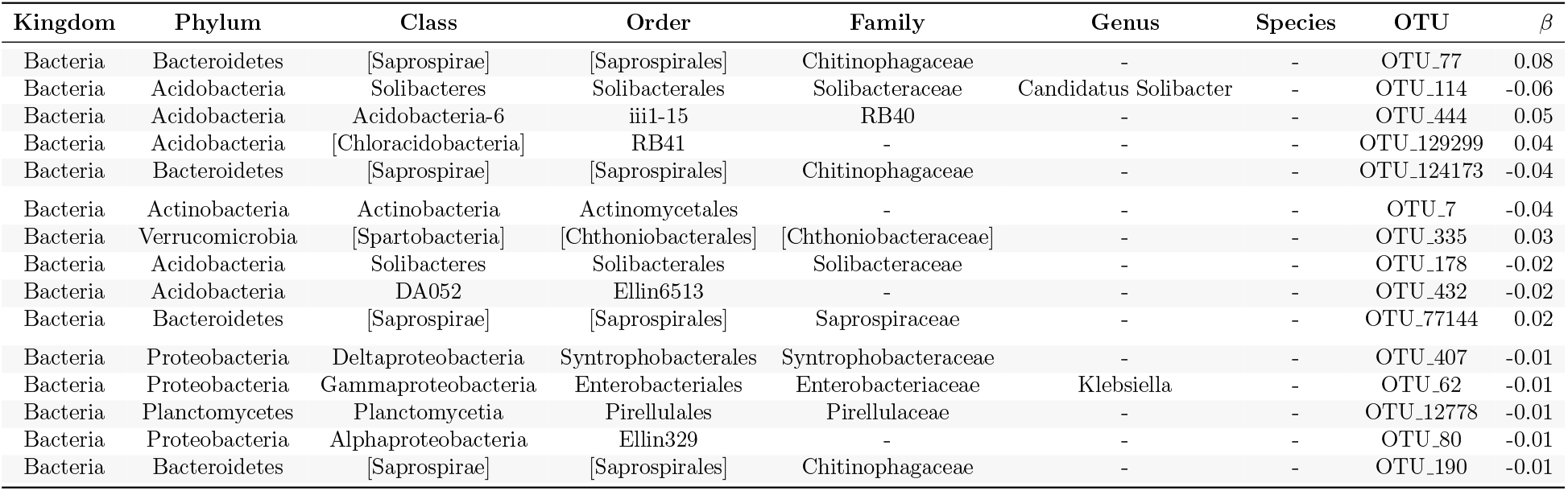
Top 15 coefficients selected by the sparse log-contrast method for Central Park Soil: pH

**Table 17:** Top 15 coefficients selected by `trac` (a = 1) for Central Park Soil: Mois

| Kingdom | Phylum | Class | Order | Family | Genus | Species | OTU | $\alpha$ |
| --- | --- | --- | --- | --- | --- | --- | --- | --- |
| Bacteria |  |  |  |  |  |  |  | -26.58 |
| Bacteria | Proteobacteria |  |  |  |  |  |  | 13.68 |
| Bacteria | Proteobacteria | Deltaproteobacteria |  |  |  |  |  | 9.71 |
| Bacteria | Proteobacteria | Alphaproteobacteria |  |  |  |  |  | 6.77 |
| Bacteria | Bacteroidetes |  |  |  |  |  |  | 4.89 |
| Bacteria | Acidobacteria |  |  |  |  |  |  | 4.68 |
| Bacteria | Bacteroidetes | Sphingobacteriia | Sphingobacteriales |  |  |  |  | -3.25 |
| Bacteria | Actinobacteria | Thermoleophilia | Gaiellales |  |  |  |  | 3.01 |
| Bacteria | Verrucomicrobia |  |  |  |  |  |  | -2.66 |
| Bacteria | Proteobacteria | Alphaproteobacteria | Sphingomonadales | Sphingomonadaceae |  |  |  | -2.40 |
| Bacteria | Actinobacteria |  |  |  |  |  |  | -2.38 |
| Bacteria | Proteobacteria | Betaproteobacteria |  |  |  |  |  | -2.16 |
| Bacteria | Acidobacteria | Acidobacteriia | Acidobacteriales | Koribacteraceae |  |  |  | -1.99 |
| Bacteria | Verrucomicrobia | [Pedosphaerae] | [Pedosphaerales] |  |  |  |  | -1.37 |
| Bacteria | Actinobacteria | Thermoleophilia | Solirubrobacterales |  |  |  |  | -1.24 |

**Table 18:**
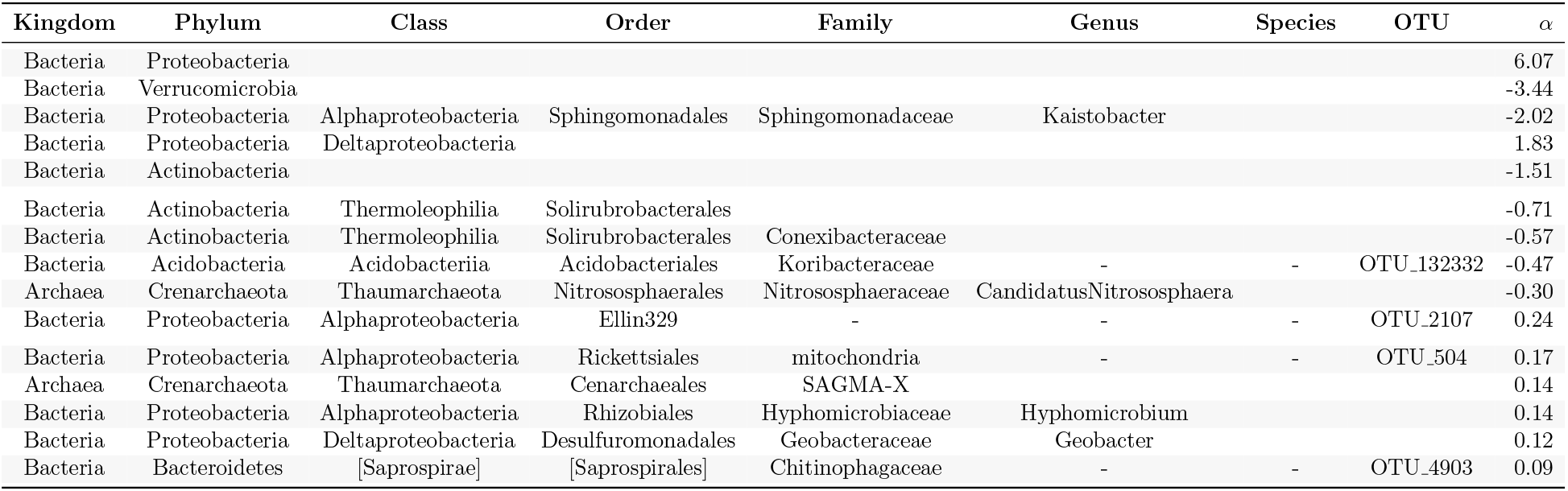
Top 15 coefficients selected by `trac` (a = 1/2) for Central Park Soil: Mois

**Table 19:** Coefficients selected by the sparse log-contrast method for Central Park Soil: Mois

| Kingdom | Phylum | Class | Order | Family | Genus | Species | OTU | $\beta$ |
| --- | --- | --- | --- | --- | --- | --- | --- | --- |
| Bacteria | Acidobacteria | Acidobacteriia | Acidobacteriales | Koribacteraceae | - | - | OTU_132332 | -0.77 |
| Bacteria | Bacteroidetes | Cytophagia | Cytophagales | Cytophagaceae | - | - | OTU_103638 | 0.46 |
| Bacteria | Bacteroidetes | [Saprospirae] | [Saprospirales] | Chitinophagaceae | - | - | OTU_4903 | 0.33 |
| Bacteria | Proteobacteria | Betaproteobacteria | MND1 | - | - | - | OTU_811 | 0.27 |
| Bacteria | Actinobacteria | Acidimicrobiia | Acidimicrobiales | - | - | - | OTU_461 | -0.20 |
| Bacteria | Proteobacteria | Gammaproteobacteria | Xanthomonadales | Sinobacteraceae | - | - | OTU_1132 | 0.20 |
| Bacteria | Planctomycetes | Phycisphaerae | WD2101 | - | - | - | OTU_132692 | -0.16 |
| Bacteria | Proteobacteria | Alphaproteobacteria | Ellin329 | - | - | - | OTU_2107 | 0.13 |
| Bacteria | Bacteroidetes | [Saprospirae] | [Saprospirales] | Chitinophagaceae | - | - | OTU_91357 | -0.12 |
| Bacteria | Proteobacteria | Alphaproteobacteria | Rickettsiales | mitochondria | - | - | OTU_504 | 0.11 |
| Bacteria | Actinobacteria | Acidimicrobiia | Acidimicrobiales | - | - | - | OTU_669 | -0.11 |
| Bacteria | Proteobacteria | Alphaproteobacteria | Sphingomonadales | Sphingomonadaceae | Kaistobacter | - | OTU_10329 | -0.09 |
| Bacteria | Actinobacteria | Acidimicrobiia | Acidimicrobiales | - | - | - | OTU_1582 | -0.08 |
| Bacteria | Verrucomicrobia | - | - | - | - | - | OTU_1207 | 0.03 |
| Archaea | Crenarchaeota | Thaumarchaeota | Cenarchaeales | SAGMA-X | - | - | OTU_208 | 0.01 |

**Table 20:** Coefficients selected by `trac` (a = 1) for Fram Strait (FL): Leucine

| Kingdom | Phylum | Class | Order | Family | Genus | Species | OTU | $\alpha$ |
| --- | --- | --- | --- | --- | --- | --- | --- | --- |
| Bacteria | Bacteroidetes | Flavobacteriia | Flavobacteriales | Flavobacteriaceae |  |  |  | 27.90 |
| Bacteria | Proteobacteria | Alphaproteobacteria |  |  |  |  |  | -23.40 |
| Bacteria | Proteobacteria | Gammaproteobacteria |  |  |  |  |  | -4.49 |

**Table 21:** Coefficients selected by `trac` (a = 1/2) for Fram Strait (FL): Leucine

| Kingdom | Phylum | Class | Order | Family | Genus | Species | OTU | $\alpha$ |
| --- | --- | --- | --- | --- | --- | --- | --- | --- |
| Bacteria | Bacteroidetes | Flavobacteriia | Flavobacteriales | Flavobacteriaceae |  |  |  | 14.30 |
| Bacteria | Proteobacteria | Alphaproteobacteria | Rhodospirillales |  |  |  |  | -8.99 |
| Bacteria | Marinimicrobia(SAR406clade) |  |  |  |  |  |  | -4.30 |
| Bacteria | Bacteroidetes | Flavobacteriia | Flavobacteriales | NS9 marine group | - | - | otu117 | -0.79 |
| Bacteria | Proteobacteria | Deltaproteobacteria | SAR324 clade(Marine group B) | - | - | - | otu14 | -0.22 |

**Table 22:** Coefficients selected by the sparse log-contrast method for Fram Strait (FL): Leucine

| Kingdom | Phylum | Class | Order | Family | Genus | Species | OTU | $\beta$ |
| --- | --- | --- | --- | --- | --- | --- | --- | --- |
| Bacteria | Bacteroidetes | Flavobacteriia | Flavobacteriales | Flavobacteriaceae | Ulvibacter | - | otu9 | 1.44 |
| Bacteria | Bacteroidetes | Flavobacteriia | Flavobacteriales | NS9 marine group | - | - | otu117 | -0.83 |
| Bacteria | Proteobacteria | Deltaproteobacteria | SAR324 clade(Marine group B) | - | - | - | otu14 | -0.61 |

**Table 23:** Coefficients selected by `trac` (a = 1) for Fram Strait (PA): Leucine

| Kingdom | Phylum | Class | Order | Family | Genus | Species | OTU | $\alpha$ |
| --- | --- | --- | --- | --- | --- | --- | --- | --- |
| Bacteria | Proteobacteria |  |  |  |  |  |  | -13.93 |
| Bacteria | Bacteroidetes | Flavobacteriia | Flavobacteriales | Flavobacteriaceae |  |  |  | 13.93 |

**Table 24:**
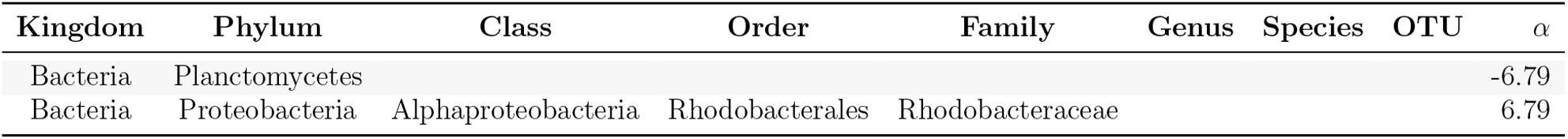
Coefficients selected by `trac` (a = 1/2) for Fram Strait (PA): Leucine

**Table 25:** Coefficients selected by the sparse log-contrast method for Fram Strait (PA): Leucine

| Kingdom | Phylum | Class | Order | Family | Genus | Species | OTU | $\beta$ |
| --- | --- | --- | --- | --- | --- | --- | --- | --- |
| Bacteria | Proteobacteria | Alphaproteobacteria | Rhodobacterales | Rhodobacteraceae | Sulfitobacter | - | otu11 | 1.41 |
| Bacteria | Proteobacteria | Deltaproteobacteria | Bdellovibrionales | Bdellovibrionaceae | OM27 clade | - | otu93 | -1.41 |

**Table 26:** Coefficients selected by `trac` (a = 1) for Ocean (TARA): Salinity

| Kingdom | Phylum | Class | Order | Family | Genus | Species | OTU | $\alpha$ |
| --- | --- | --- | --- | --- | --- | --- | --- | --- |
| Bacteria | Proteobacteria | Alphaproteobacteria |  |  |  |  |  | 4.00 |
| Bacteria |  |  |  |  |  |  |  | -2.92 |
| Bacteria | Bacteroidetes |  |  |  |  |  |  | -1.38 |
| Bacteria | Proteobacteria | Gammaproteobacteria |  |  |  |  |  | 0.30 |

**Table 27:** Coefficients selected by `trac` (a = 1/2) for Ocean (TARA): Salinity

| Kingdom | Phylum | Class | Order | Family | Genus | Species | OTU | $\alpha$ |
| --- | --- | --- | --- | --- | --- | --- | --- | --- |
| Bacteria | Bacteroidetes | Flavobacteria | Flavobacteriales | NS9marinegroup |  |  |  | -0.96 |
| Bacteria | Proteobacteria | Alphaproteobacteria | SAR11clade |  |  |  |  | 0.55 |
| Bacteria | Proteobacteria | Alphaproteobacteria |  |  |  |  |  | 0.38 |
| Bacteria | Proteobacteria | Gammaproteobacteria | Oceanospirillales | Halomonadaceae |  |  |  | -0.37 |
| Bacteria | Cyanobacteria |  |  |  |  |  |  | 0.25 |
| Bacteria | Cyanobacteria | Cyanobacteria | SubsectionI | FamilyI | Synechococcus |  |  | 0.12 |
| Bacteria | Proteobacteria | Gammaproteobacteria | E01-9C-26 marine group | - | - | JF747664.1.1516 | OTU520 | 0.09 |
| Bacteria | Proteobacteria | Gammaproteobacteria | Alteromonadales | Alteromonadaceae | Marinobacter |  |  | -0.07 |
| Bacteria | Cyanobacteria | Cyanobacteria |  |  |  |  |  | 0.02 |
| Bacteria | Bacteroidetes |  |  |  |  |  |  | -0.02 |

**Table 28:** Coefficients selected by the sparse log-contrast method for Ocean (TARA): Salinity

| Kingdom | Phylum | Class | Order | Family | Genus | Species | OTU | $\beta$ |
| --- | --- | --- | --- | --- | --- | --- | --- | --- |
| Bacteria | Proteobacteria | Gammaproteobacteria | E01-9C-26 marine group | - | - | JF747664.1.1516 | OTU520 | 0.25 |
| Bacteria | Proteobacteria | Gammaproteobacteria | Oceanospirillales | JL-ETNP-Y6 | - | GQ347814.1.1378 | OTU925 | -0.11 |
| Bacteria | Proteobacteria | Gammaproteobacteria | Oceanospirillales | SAR86 clade | - | AACY020549891.3846.5359 | OTU19 | 0.06 |
| Bacteria | Verrucomicrobia | Verrucomicrobiae | Verrucomicrobiales | Verrucomicrobiaceae | Roseibacillus | GU062019.1.1504 | OTU729 | -0.05 |
| Bacteria | Proteobacteria | Gammaproteobacteria | Alteromonadales | Alteromonadaceae | Melitea | HQ326447.1.1497 | OTU2376 | -0.05 |
| Bacteria | Proteobacteria | Gammaproteobacteria | Salinisphaerales | Salinisphaeraceae | Salinisphaera | AB735546.1.1462 | OTU1096 | -0.04 |
| Bacteria | Bacteroidetes | Flavobacteria | Flavobacteriales | NS9 marine group | - | HQ673682.1.1487 | OTU1168 | -0.04 |
| Bacteria | Proteobacteria | Gammaproteobacteria | Alteromonadales | Idiomarinaceae | Idiomarina | EU440983.1.1508 | OTU2517 | -0.02 |
| Bacteria | Actinobacteria | Acidimicrobiia | Acidimicrobiales | OCS155 marine group | - | AACY020396101.1882.3388 | OTU56 | 0.01 |

Figure 3 compares the aggregations across the taxonomic tree that were found by standard `trac` for soil pH and moisture prediction, respectively. We observe that only the phyla Bacteroidetes and Verrucomicrobia, and the order Acidobacteriales are common in both models, confirming that the relevant taxonomic aggregations depend on the response variable being predicted.

Finally, we observe similar prediction performance in terms of test error (40 – 45), with standard `trac` being outperformed by the other methods across all base level aggregations. For moisture prediction, weighted `trac` provides an excellent trade-off between model inter-pretability and predictability.

**Table 2:** Average out-of-sample test errors (model sparsity in parenthesis) for `trac` (*a* = {1, 1/2}) and sparse log-contrast models, respectively. Each row considers a different base level (OTU, genus, and family). Each number is averaged over ten different training/test splits of the Central Park soil, Moisture data.

| Base Level | $p$ | **trac** ($a = 1$) | **trac** ($a = 1/2$) | Sparse Log-Contrast |
| --- | --- | --- | --- | --- |
| OTU | 3379 | 42 (8) | 40 (13) | 40 (23) |
| Genus | 2779 | 42 (5) | 40 (17) | 41 (19) |
| Family | 1492 | 45 (4) | 42 (12) | 41 (16) |

#### Primary bacterial production in the Fram Strait

Current estimates suggest that the ocean microbiome could be responsible for about half of all primary production occurring on Earth [13, 14]. While net primary production is known to be highly influenced by a multitude of environmental drivers, including light, nutrients, and temperature [15], it is not yet established whether amplicon sequencing data alone contain enough information to serve as a stable predictor of (regional) marine primary production.

To investigate this relationship we consider a marine dataset, put forward in [16], that covers the Fram Strait, the main gateway between the North Atlantic and Arctic Oceans. The Fram Strait comprises two distinct oceanic regions, the northward flowing West Spits-bergen Current (WSC), and the East Greenland Current (EGC) flowing southward along the Greenland shelf. Recent ocean simulations, however, suggest substantial horizontal mixing and exchange by eddies between the two regions. We thus trained regression models from amplicon data across both regions and considered the available leucine incorporation (as proxy to bacterial production) as the outcome [16]. We learned separate models for the two different size fractions: *p* = 4530 free-living (FL) taxa in the 0.22*μm* fraction, and *p* = 3320 particle-associated (PA) taxa in 3*μm* fraction.

**Figure 4:**
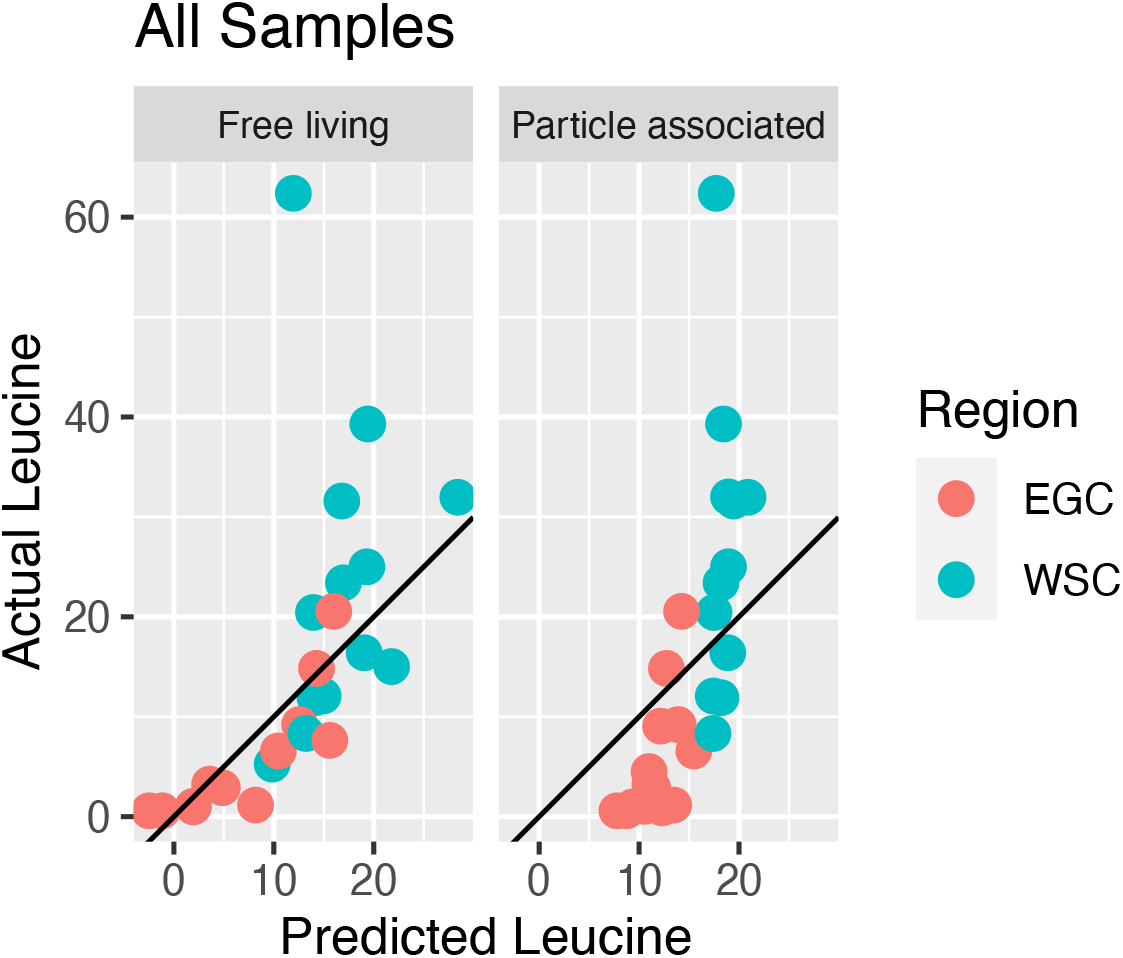
Predictions by `trac` (*a* = 1) of primary production (leucine) from free living (FL) and particle associated (PA) taxa. The data points are colored by region in the Fram Strait: West Spitsbergen Current (WSC), and the East Greenland Current (EGC). The correlation between predicted and measured leucine (on the test set of split 1) is 0.57 for FL taxa and 0.90 and PA taxa, respectively. Tables 20 and 23 show the selected taxa for these models.

On the FL dataset, `trac` (*a* = 1) identifies a parsimonious model, comprising three aggregated taxonomic groups, strongly associated with bacterial production. The two classes Gammaproteobacteria and Alphaproteobacteria are negatively associated, and the family Flavobacteriaceae is positively associated with bacterial production, leading to a two-factor log-contrast model. On the PA dataset, standard `trac` infers a single predictive log-contrast with the Flavobacteriaceae family being positively associated and the entire phylum Proteobacteria negatively associated with primary production. On the test data (split 1), the PA model predictions show a correlation of 0.90 with the measurements. Figure 4 summarizes the scatter plots of leucine measurements vs. `trac` predictions for the two size fractions, colored by region WSC and EGC, respectively.

We observe that the PA model appears to serve as an implicit region classifier since predicted leucine values of < 17 belong uniquely to samples in the low-productivity EGC region (see top right panel in Figure 4). Our model suggests an important positive association of the heterotrophic Flavobacteriaceae with primary production, independent of size class. Flavobacteriaceae are known to strongly contribute to mineralization of primary-produced organic matter (see [17] and references therein), thus suggesting an indirect relationship between Flavobacteriaceae and primary production. However, previous studies in South polar front and antarctic zone postulated a strong role of Flavobacteriaceae for polar primary production [18].

As highlighted in Tables 3 and 4, weighted `trac` and log-contrast models lead to sparse models and outperform standard `trac` in terms of average test error. In the FL data set (data split 1), weighted `trac` selects both higher order aggregations and two OTUs both of which are also selected by the log-contrast models. For the PA dataset, all models result in single log-ratio models, either on the phylum/family level or OTU level, respectively.

**Table 3:** Average out-of-sample test errors (model sparsity in parenthesis) for `trac` (*a* = {1, 1/2}) and sparse log-contrast models, respectively. Each row considers a different base level (OTU, genus, and family). Each number is averaged over ten different training/test splits of the Fram Strait (PA) data.

| Base Level | $p$ | **trac** ($a = 1$) | **trac** ($a = 1/2$) | Sparse Log-Contrast |
| --- | --- | --- | --- | --- |
| OTU | 3320 | 1.3e+02 (4) | 1.2e+02 (5) | 84 (5) |
| Genus | 1796 | 1.1e+02 (5) | 1e+02 (4) | 81 (4) |
| Family | 597 | 1.2e+02 (3) | 1e+02 (4) | 99 (6) |

#### Global predictive model of ocean salinity from Tara data

We complement the Tara data set analysis from the main text with showing the scatter plot of measured vs. predicted salinity for the standard `trac` model (trained on data split 1) in Figure 5.

**Figure 5:**
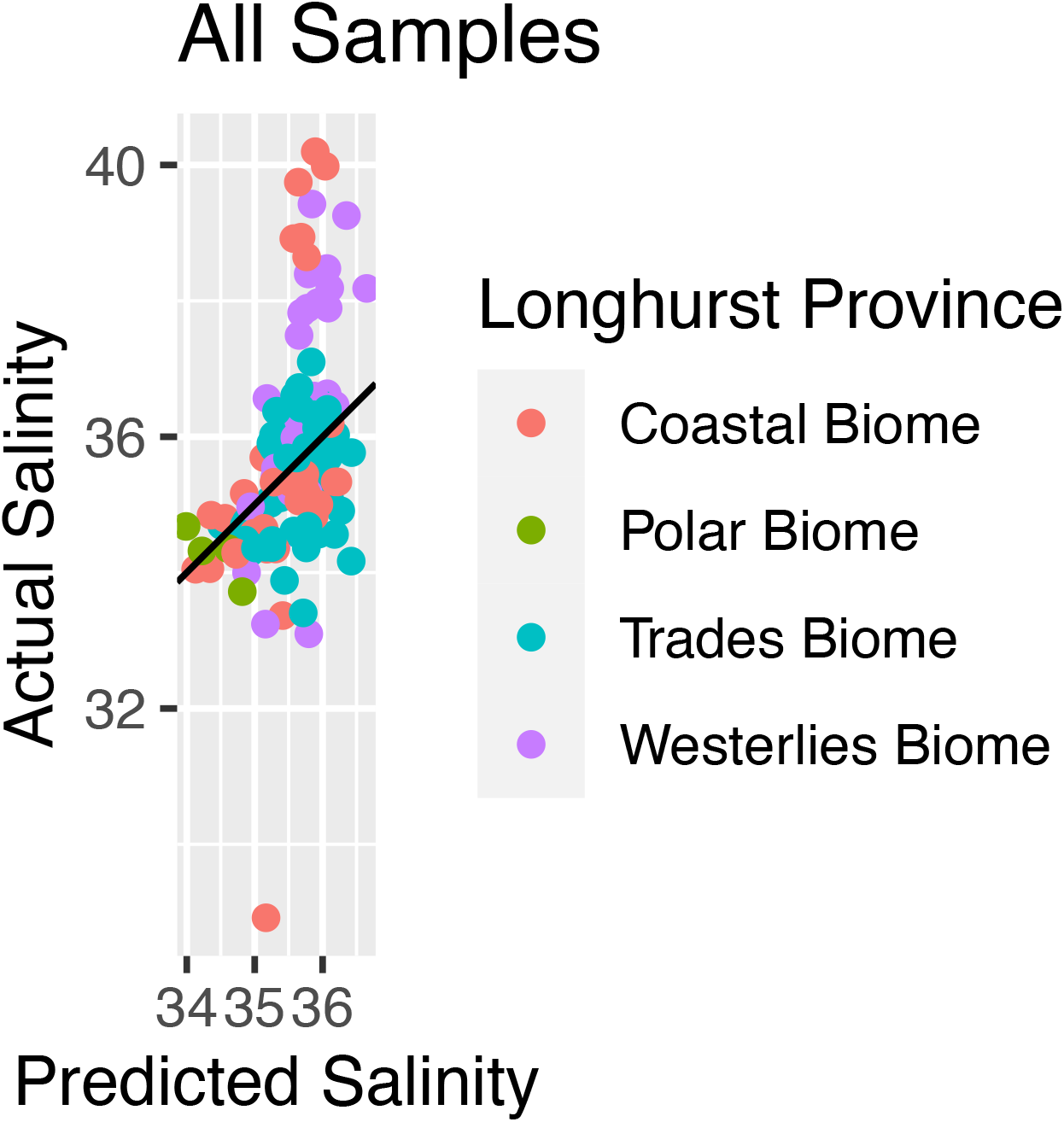
Measured salinity (y-axis) vs. standard `trac` (*a* =1) model prediction (x-axis) on the Tara data (model training performed on data split 1). Each sample is colored by one of the four Longhurst Biome definitions. Outliers to the model are located in Coastal and Westerlies Biomes.

### D Additional Selected Coefficient Tables

## References

[1] Ron Sender, Shai Fuchs, and Ron Milo. Revised Estimates for the Number of Human and Bacteria Cells in the Body. PLoS Biology, 14(8):1–14, 2016.

[2] Yinon M. Bar-On, Rob Phillips, and Ron Milo. The biomass distribution on Earth. Proceedings of the National Academy of Sciences of the United States of America, 115(25):6506–6511, 2018.

[3] Shinichi Sunagawa, Luis Pedro Coelho, Samuel Chaffron, Jens Roat Kultima, Karine Labadie, Guillem Salazar, Bardya Djahanschiri, Georg Zeller, Daniel R. Mende, Adriana Alberti, Francisco M. Cornejo-Castillo, Paul I. Costea, Corinne Cruaud, Francesco D’Ovidio, Stefan Engelen, Isabel Ferrera, Josep M. Gasol, Lionel Guidi, Falk Hildebrand, Florian Kokoszka, Cyrille Lepoivre, Gipsi Lima-Mendez, Julie Poulain, Bonnie T. Poulos, Marta Royo-Llonch, Hugo Sarmento, Sara Vieira-Silva, Céline Dimier, Marc Picheral, Sarah Searson, Stefanie Kandels-Lewis, Emmanuel Boss, Michael Follows, Lee Karp-Boss, Uros Krzic, Emmanuel G. Reynaud, Christian Sardet, Mike Sieracki, Didier Velayoudon, Chris Bowler, Colomban De Vargas, Gabriel Gorsky, Nigel Grimsley, Pascal Hingamp, Daniele Iudicone, Olivier Jaillon, Fabrice Not, Hiroyuki Ogata, Stephane Pesant, Sabrina Speich, Lars Stemmann, Matthew B. Sullivan, Jean Weissenbach, Patrick Wincker, Eric Karsenti, Jeroen Raes, Silvia G. Acinas, and Peer Bork. Structure and function of the global ocean microbiome. Science, 348(6237):1–10, 2015.

[4] Mohammad Bahram, Falk Hildebrand, Sofia K Forslund, Jennifer L Anderson, Nadejda A Soudzilovskaia, Peter M Bodegom, Johan Bengtsson-Palme, Sten Anslan, Luis Pedro Coelho, Helery Harend, Jaime Huerta-Cepas, Marnix H Medema, Mia R Maltz, Sunil Mundra, Pål Axel Olsson, Mari Pent, Sergei Põlme, Shinichi Sunagawa, Martin Ryberg, Leho Tedersoo, and Peer Bork. Structure and function of the global topsoil microbiome. Nature, 560(7717):233–237, 2018.

[5] Daniel et al. McDonald. American gut: an open platform for citizen science microbiome research. mSystems, 3(3), 2018.

[6] Benjamin J. Callahan, Paul J. McMurdie, and Susan P. Holmes. Exact sequence variants should replace operational taxonomic units in marker-gene data analysis. ISME Journal, 11(12):2639–2643, 2017.

[7] Q. Wang, G. M. Garrity, J. M. Tiedje, and J. R. Cole. Naive Bayesian classifier for rapid assignment of rRNA sequences into the new bacterial taxonomy. Appl. Environ. Microbiol., 73(16):5261–5267, Aug 2007.

[8] Daniel McDonald, Morgan N Price, Julia Goodrich, Eric P Nawrocki, Todd Z DeSantis, Alexander Probst, Gary L Andersen, Rob Knight, and Philip Hugenholtz. An improved Greengenes taxonomy with explicit ranks for ecological and evolutionary analyses of bacteria and archaea. The ISME Journal, 6(3):610–618, 2012.

[9] Christian Quast, Elmar Pruesse, Pelin Yilmaz, Jan Gerken, Timmy Schweer, Pablo Yarza, Jörg Peplies, and Frank Oliver Glöckner. The SILVA ribosomal RNA gene database project: Improved data processing and web-based tools. Nucleic Acids Research, 41(D1):590–596, 2013.

[10] N. Chaudhary, A. K. Sharma, P. Agarwal, A. Gupta, and V. K. Sharma. 16S classifier: a tool for fast and accurate taxonomic classification of 16S rRNA hypervariable regions in metagenomic datasets. PLoS ONE, 10(2):e0116106, 2015.

[11] Klaus Peter Schliep. phangorn: Phylogenetic analysis in R. Bioinformatics, 27(4):592–593, 2011.

[12] Tong Zhang, Ming-Fei Shao, and Lin Ye. 454 pyrosequencing reveals bacterial diversity of activated sludge from 14 sewage treatment plants. The ISME Journal, 6(6):1137–1147, 2012.

[13] Jun Chen, Frederic D. Bushman, James D. Lewis, Gary D. Wu, and Hongzhe Li. Structure-constrained sparse canonical correlation analysis with an application to microbiome data analysis. Biostatistics, 14(2):244–258, 2013.

[14] Fan Xia, Jun Chen, Wing Kam Fung, and Hongzhe Li. A logistic normal multinomial regression model for microbiome compositional data analysis. Biometrics, 69(4):1053–1063, 2013.

[15] Wei Lin, Pixu Shi, Rui Feng, and Hongzhe Li. Variable selection in regression with compositional covariates. Biometrika, 101:785–797, 11 2014.

[16] T. W. Randolph, S. Zhao, W. Copeland, M. Hullar, and A. Shojaie. Kernel-Penalized Regression for Analysis of Microbiome Data. ArXiv e-prints, November 2015.

[17] J. Aitchison. The statistical analysis of compositional data. Journal of the Royal Statistical Society. Series B (Methodological), 44(2):139–177, 1982.

[18] Juan Jose Egozcue and Vera Pawlowsky-Glahn. Groups of parts and their balances in compositional data analysis. Mathematical Geology, 37(7):795–828, 2005.

[19] Gregory B. Gloor, Jean M. Macklaim, Vera Pawlowsky-Glahn, and Juan J. Egozcue. Microbiome Datasets Are Compositional: And This Is Not Optional. Frontiers in Microbiology, 8(November):2224, 2017.

[20] J Bacon-Shone and J Aitchison. Log contrast models for experiments with mixtures. Biometrika, 1984.

[21] Xiaohan Yan and Jacob Bien. Rare feature selection in high dimensions. Journal of the American Statistical Association, 0(just-accepted):1–30, 2020.

[22] Catherine Lozupone and Rob Knight. UniFrac: a New Phylogenetic Method for Comparing Microbial Communities UniFrac: a New Phylogenetic Method for Comparing Microbial Communities. Applied and environmental microbiology, 71(12):8228–8235, 2005.

[23] Alex D. Washburne, Justin D. Silverman, Jonathan W. Leff, Dominic J. Bennett, John L. Darcy, Sayan Mukherjee, Noah Fierer, and Lawrence A. David. Phylogenetic factorization of compositional data yields lineage-level associations in microbiome datasets. PeerJ, 5:e2969, 2017.

[24] Justin D Silverman, Alex D Washburne, Sayan Mukherjee, and Lawrence A David. A phylogenetic transform enhances analysis of compositional microbiota data. eLife, 6:1–20, 2017.

[25] James T. Morton, Jon Sanders, Robert A. Quinn, Daniel McDonald, Antonio Gonzalez, Yoshiki Vázquez-Baeza, Jose A. Navas-Molina, Se Jin Song, Jessica L. Metcalf, Embriette R. Hyde, Manuel Lladser, Pieter C. Dorrestein, and Rob Knight. Balance Trees Reveal Microbial Niche Differentiation. mSystems, 2(1):e00162—16, 2017.

[26] Alex D. Washburne, Justin D. Silverman, James T. Morton, Daniel J. Becker, Daniel Crowley, Sayan Mukherjee, Lawrence A. David, and Raina K. Plowright. Phylofactorization: a graph partitioning algorithm to identify phylogenetic scales of ecological data. Ecological Monographs, 89(2):1–27, 2019.

[27] J. Zhai, J. Kim, K. S. Knox, H. L. Twigg, H. Zhou, and J. J. Zhou. Variance Component Selection With Applications to Microbiome Taxonomic Data. Front Microbiol, 9:509, 2018.

[28] Jian Xiao, Li Chen, Stephen Johnson, Yue Yu, Xianyang Zhang, and Jun Chen. Predictive modeling of microbiome data using a phylogeny-regularized generalized linear mixed model. Frontiers in Microbiology, 9(JUN):1–14, 2018.

[29] M. Khabbazian, R. Kriebel, K. Rohe, and C. Ané. Fast and accurate detection of evolutionary shifts in ornstein–uhlenbeck models. Methods in Ecology and Evolution, 7(7):811–824, 2016.

[30] Tao Wang and Hongyu Zhao. Structured subcomposition selection in regression and its application to microbiome data analysis. The Annals of Applied Statistics, 11(2):771—791, 2017.

[31] Patrick H. Bradley, Stephen Nayfach, and Katherine S. Pollard. Phylogeny-corrected identification of microbial gene families relevant to human gut colonization. PLoS Computational Biology, 14(8):1–41, 2018.

[32] Robert Tibshirani. Regression shrinkage and selection via the lasso. Journal of the Royal Statistical Society, Series B, 58:267–288, 1996.

[33] Patrick L Combettes and Christian L Müller. Regression models for compositional data: General log-contrast formulations, proximal optimization, and microbiome data applications. Statistics in Biosciences, pages 1–26, 2020.

[34] Brian R. Gaines, Juhyun Kim, and Hua Zhou. Algorithms for Fitting the Constrained Lasso. Journal of Computational and Graphical Statistics, 27(4):861–871, 2018.

[35] Léo Simpson, Patrick L. Combettes, and Christian L Müller. c-lasso - a Python package for constrained sparse and robust regression and classification. Journal of Open Source Software, 6(57):2844, 2021.

[36] Kevin Ushey, JJ Allaire, and Yuan Tang. reticulate: Interface to ‘Python’, 2020. R package version 1.16.

[37] Paul J. McMurdie and Susan Holmes. phyloseq: An R Package for Reproducible Interactive Analysis and Graphics of Microbiome Census Data. PLoS ONE, 8(4):e61217, 2013.

[38] Hadley Wickham. ggplot2: Elegant Graphics for Data Analysis. Springer-Verlag New York, 2016.

[39] E. Paradis and K. Schliep. ape 5.0: an environment for modern phylogenetics and evolutionary analyses in R. Bioinformatics, 35:526–528, 2019.

[40] Gabor Csardi and Tamas Nepusz. The igraph software package for complex network research. Inter Journal, Complex Systems:1695, 2006.

[41] Guangchuang Yu, David K Smith, Huachen Zhu, Yi Guan, and Tommy Tsan-Yuk Lam. ggtree: an r package for visualization and annotation of phylogenetic trees with their covariates and other associated data. Methods in Ecology and Evolution, 8(1):28–36, 2017.

[42] Trevor Hastie, Robert Tibshirani, and Jerome Friedman. The elements of statistical learning: data mining, inference, and prediction. Springer Science & Business Media, 2009.

[43] J. Rivera-Pinto, J. J. Egozcue, V. Pawlowsky-Glahn, R. Paredes, M. Noguera-Julian, and M. L. Calle. Balances: a New Perspective for Microbiome Analysis. mSystems, 3(4):1–12, 2018.

[44] Michelle Badri, Zachary D Kurtz, Richard Bonneau, and Christian L Müller. Shrinkage improves estimation of microbial associations under different normalization methods. bioRxiv, 2020.

[45] Kelly S Ramirez, Jonathan W Leff, Albert Barberán, Scott Thomas Bates, Jason Betley, Thomas W Crowther, Eugene F Kelly, Emily E Oldfield, E. Ashley Shaw, Christopher Steenbock, Mark A Bradford, Diana H Wall, and Noah Fierer. Biogeographic patterns in below-ground diversity in New York City’s Central Park are similar to those observed globally. Proceedings of the Royal Society B: Biological Sciences, 281(1795), 2014.

[46] Eduard Fadeev, Ian Salter, Vibe Schourup-Kristensen, Eva Maria Nöthig, Katja Metfies, Anja Engel, Judith Piontek, Antje Boetius, and Christina Bienhold. Microbial communities in the east and west fram strait during sea ice melting season. Frontiers in Marine Science, 5(NOV):1–21, 2018.

[47] Stephanie M. Dillon, Daniel N. Frank, and Cara C. Wilson. The gut microbiome and HIV-1 pathogenesis: A two-way street. Aids, 30(18):2737–2751, 2016.

[48] Piotr Nowak, Marius Troseid, Ekatarina Avershina, Babilonia Barqasho, Ujjwal Neogi, Kristian Holm, Johannes R. Hov, Kajsa Noyan, Jan Vesterbacka, Jenny Svärd, Knut Rudi, and Anders Sönnerborg. Gut microbiota diversity predicts immune status in HIV-1 infection. Aids, 29(18):2409–2418, 2015.

[49] Netanya G. Sandler, Handan Wand, Annelys Roque, Matthew Law, Martha C. Nason, Daniel E. Nixon, Court Pedersen, Kiat Ruxrungtham, Sharon R. Lewin, Sean Emery, James D. Neaton, Jason M. Brenchley, Steven G. Deeks, Irini Sereti, and Daniel C. Douek. Plasma levels of soluble CD14 independently predict mortality in HIV infection. Journal of Infectious Diseases, 203(6):780–790, 2011.

[50] Grégory Dubourg. Impact of HIV on the human gut microbiota: Challenges and perspectives. Human Microbiome Journal, 2:3–9, 2016.

[51] Cynthia L. Monaco, David B Gootenberg, Guoyan Zhao, Scott A Handley, Musie S Ghebremichael, Efrem S Lim, Alex Lankowski, Megan T. Baldridge, Craig B. Wilen, Meaghan Flagg, Jason M. Norman, Brian C. Keller, Jesús Mario Luévano, David Wang, Yap Boum, Jeffrey N. Martin, Peter W. Hunt, David R. Bangsberg, Mark J Siedner, Douglas S Kwon, and Herbert W Virgin. Altered Virome and Bacterial Microbiome in Human Immunodeficiency Virus-Associated Acquired Immunodeficiency Syndrome. Cell Host and Microbe, 19(3):311–322, 2016.

[52] Noah Fierer and Robert B Jackson. The diversity and biogeography of soil bacterial communities. PNAS, 103(3), 2006.

[53] Christian L Lauber, Micah Hamady, Rob Knight, and Noah Fierer. Pyrosequencing-Based Assessment of Soil pH as a Predictor of Soil Bacterial Community Structure at the Continental Scale. Applied and Environmental Microbiology, 75(15):5111–5120, 2009.

[54] Andrea K. Bartram, Xingpeng Jiang, Michael D.J. Lynch, Andre P. Masella, Graeme W. Nicol, Jonathan Dushoff, and Josh D. Neufeld. Exploring links between pH and bacterial community composition in soils from the Craibstone Experimental Farm. FEMS Microbiology Ecology, 87(2):403–415, 2014.

[55] Shinichi Sunagawa, Silvia G. Acinas, Peer Bork, Chris Bowler, Silvia G. Acinas, Marcel Babin, Peer Bork, Emmanuel Boss, Chris Bowler, Guy Cochrane, Colomban de Vargas, Michael Follows, Gabriel Gorsky, Nigel Grimsley, Lionel Guidi, Pascal Hingamp, Daniele Iudicone, Olivier Jaillon, Stefanie Kandels, Lee Karp-Boss, Eric Karsenti, Magali Lescot, Fabrice Not, Hiroyuki Ogata, Stéphane Pesant, Nicole Poulton, Jeroen Raes, Christian Sardet, Mike Sieracki, Sabrina Speich, Lars Stemmann, Matthew B. Sullivan, Shinichi Sunagawa, Patrick Wincker, Damien Eveillard, Gabriel Gorsky, Lionel Guidi, Daniele Iudicone, Eric Karsenti, Fabien Lombard, Hiroyuki Ogata, Stephane Pesant, Matthew B. Sullivan, Patrick Wincker, and Colomban de Vargas. Tara Oceans: towards global ocean ecosystems biology. Nature Reviews Microbiology, 18(8):428–445, 2020.

[56] Ramiro Logares, Shinichi Sunagawa, Guillem Salazar, Francisco M. Cornejo-Castillo, Isabel Ferrera, Hugo Sarmento, Pascal Hingamp, Hiroyuki Ogata, Colomban de Vargas, Gipsi Lima-Mendez, Jeroen Raes, Julie Poulain, Olivier Jaillon, Patrick Wincker, Stefanie Kandels-Lewis, Eric Karsenti, Peer Bork, and Silvia G. Acinas. Metagenomic 16S rDNA Illumina tags are a powerful alternative to amplicon sequencing to explore diversity and structure of microbial communities. Environmental Microbiology, 2014.

[57] Thierry C. Bouvier and Paul A. Del Giorgio. Compositional changes in free-living bacterial communities along a salinity gradient in two temperate estuaries. Limnology and Oceanography, 47(2):453–470, 2002.

[58] Matthew T. Cottrell and David L. Kirchman. Contribution of major bacterial groups to bacterial biomass production (thymidine and leucine incorporation) in the Delaware estuary. Limnology and Oceanography, 48(1 I):168–178, 2003.

[59] Pelin Yilmaz, Pablo Yarza, Josephine Z. Rapp, and Frank O. Glöckner. Expanding the world of marine bacterial and archaeal clades. Frontiers in Microbiology, 6(JAN):1–29, 2016.

[60] Pixu Shi, Anru Zhang, and Hongzhe Li. Regression analysis for microbiome compositional data. Ann. Appl. Stat., 10(2):1019–1040, 06 2016.

[61] Ruth E Ley, Fredrik Bäckhed, Peter Turnbaugh, Catherine A Lozupone, Robin D Knight, and Jeffrey I Gordon. Obesity alters gut microbial ecology. Proceedings of the National Academy of Sciences of the United States of America, 102(31):11070–11075, 2005.

[62] P. J. Turnbaugh, M. Hamady, T. Yatsunenko, B. L. Cantarel, A. Duncan, R. E. Ley, M. L. Sogin, W. J. Jones, B. A. Roe, J. P. Affourtit, M. Egholm, B. Henrissat, A. C. Heath, R. Knight, and J. I. Gordon. A core gut microbiome in obese and lean twins. Nature, 457(7228):480–484, Jan 2009.

[63] Antoine Bichat, Jonathan Plassais, Christophe Ambroise, and Mahendra Mariadassou. Incorporating Phylogenetic Information in Microbiome Differential Abundance Studies Has No Effect on Detection Power and FDR Control. Frontiers in Microbiology, 11(April):1–13, 2020.

[64] Aditya Mishra and Christian L. Müller. Robust regression with compositional covariates. arXiv preprint arXiv:1909.04990, 2019.

[65] Saharon Rosset and Ji Zhu. Piecewise linear regularized solution paths. Annals of Statistics, 35(3):1012–1030, 2007.

[66] Xiaohan Yan. Statistical Learning for Structural Patterns with Trees. PhD thesis, Cornell University, 2018.

## References

[1] Léo Simpson, Patrick L. Combettes, and Christian L Müller. c-lasso - a Python package for constrained sparse and robust regression and classification. Journal of Open Source Software, 6(57):2844, 2021.

[2] Xiaohan Yan and Jacob Bien. Rare feature selection in high dimensions. Journal of the American Statistical Association, 0(just-accepted):1–30, 2020.

[3] P. J. Turnbaugh, M. Hamady, T. Yatsunenko, B. L. Cantarel, A. Duncan, R. E. Ley, M. L. Sogin, W. J. Jones, B. A. Roe, J. P. Affourtit, M. Egholm, B. Henrissat, A. C. Heath, R. Knight, and J. I. Gordon. A core gut microbiome in obese and lean twins. Nature, 457(7228):480–484, Jan 2009.

[4] Ruth E. Ley, Peter J. Turnbaugh, Samuel Klein, and Jeffrey I. Gordon. Microbial ecology: Human gut microbes associated with obesity. Nature, 2006.

[5] Wei Lin, Pixu Shi, Rui Feng, and Hongzhe Li. Variable selection in regression with compositional covariates. Biometrika, 101:785–797, 11 2014.

[6] Pixu Shi, Anru Zhang, and Hongzhe Li. Regression analysis for microbiome compositional data. Ann. Appl. Stat., 10(2):1019–1040, 06 2016.

[7] G. D. Wu, J. Chen, C. Hoffmann, K. Bittinger, Y.-Y. Chen, S. A. Keilbaugh, M. Bewtra, D. Knights, W. A. Walters, R. Knight, R. Sinha, E. Gilroy, K. Gupta, R. Baldassano, L. Nessel, H. Li, F. D. Bushman, and J. D. Lewis. Linking Long-Term Dietary Patterns with Gut Microbial Enterotypes. Science, 334(6052):105–108, 2011.

[8] Kaihei Oki, Mutsumi Toyama, Taihei Banno, Osamu Chonan, Yoshimi Benno, and Koichi Watanabe. Comprehensive analysis of the fecal microbiota of healthy Japanese adults reveals a new bacterial lineage associated with a phenotype characterized by a high frequency of bowel movements and a lean body type. BMC Microbiology, pages 5–11, 2016.

[9] Alejandra Chávez-Carbajal, Khemlal Nirmalkar, Ana Pérez-Lizaur, Fernando Hernández-Quiroz, Silvia Ramírez-Del-Alto, Jaime García-Mena, and César Hernández-Guerrero. Gut microbiota and predicted metabolic pathways in a sample of Mexican women affected by obesity and obesity plus metabolic syndrome. International Journal of Molecular Sciences, 20(2):1–18, 2019.

[10] Noah Fierer and Robert B Jackson. The diversity and biogeography of soil bacterial communities. PNAS, 103(3), 2006.

[11] Quanchao Zeng, Yanghong Dong, and Shaoshan An. Bacterial community responses to soils along a latitudinal and vegetation gradient on the Loess Plateau, China. PLoS ONE, 11(4):1–17, 2016.

[12] Alexandre B. de Menezes, Christoph Müller, Nicholas Clipson, and Evelyn Doyle. The soil microbiome at the Gi-FACE experiment responds to a moisture gradient but not to CO2 enrichment. Microbiology (United Kingdom), 162(9):1572–1582, 2016.

[13] Alan Longhurst, Shubha Sathyendranath, Trevor Platt, and Carla Caverhill. An estimate of global primary production in the ocean from satellite radiometer data. Journal of Plankton Research, 17(6):1245–1271, 1995.

[14] Mary Ann Moran. The global ocean microbiome. Science, 350(6266), 2015.

[15] P W Boyd, S Sundby, and H.-O. Pörtner. Net primary production in the ocean. Climate Change 2014: Impacts, Adaptation, and Vulnerability. Part A: Global and Sectoral Aspects. Contribution of Working Group II to the Fifth Assessment Report of the Intergovernmental Panel on Climate Change, pages 133–136, 2014.

[16] Eduard Fadeev, Ian Salter, Vibe Schourup-Kristensen, Eva Maria Nöthig, Katja Metfies, Anja Engel, Judith Piontek, Antje Boetius, and Christina Bienhold. Microbial communities in the east and west fram strait during sea ice melting season. Frontiers in Marine Science, 5(NOV):1–21, 2018.

[17] John P. Bowman and David S. Nichols. Novel members of the family Flavobacteriaceae from Antarctic maritime habitats including Subsaximicrobium wynnwilliamsii gen. nov., sp. nov., Subsaximicrobium saxinquilinus sp. nov., Subsaxibacter broadyi gen. nov., sp. nov., Lacinutrix copepodicola gen. nov., sp. nov., and novel species of the genera Bizionia, Gelidibacter and Gillisia. International Journal of Systematic and Evolutionary Microbiology, 55(4):1471–1486, 2005.

[18] Guy C.J. Abell and John P. Bowman. Ecological and biogeographic relationships of class Flavobacteria in the Southern Ocean. FEMS Microbiology Ecology, 51(2):265–277, 2005.

